# The evolution of public goods altruism

**DOI:** 10.64898/2025.12.08.693068

**Authors:** Florian J. F. Labourel, Philip G. Madgwick, Christopher R. L. Thompson, Jason B. Wolf

## Abstract

Organisms often behave altruistically by investing resources into public goods production. Because the rewards of public goods are freely available to everyone in a group, they are potentially vulnerable to exploitation by cheaters. Classic kin selection models offer solutions to this public goods dilemma in terms of *why* public goods investment can be favoured, while game theoretical models offer insights into the conditions where exploitative cheaters can persist. However, existing theory does not provide a unified framework for understanding how much individuals should actually invest in altruism or how this investment strategy relates to why cheaters can coexist with altruists. Here we fill this gap by considering how resource allocation trade-offs shape the generosity of altruists, how this investment decision, in turn, sets the conditions for cheater coexistence, and how local competition impacts these outcomes. We find that, whenever altruists can evolve the optimal level of public goods investment, they exclude cheaters, revealing that stable altruists/cheater coexistence requires overly generous altruists—an outcome frequently driven by mechanistic constraints that limit the range of behaviours in biological systems. Importantly, we show that local competition tends to prevent such coexistence. By capturing key biologically relevant factors, our model provides a compelling framework for interpreting patterns of variation in altruistic behaviour in natural populations, and a means of generating testable predictions.

## Introduction

Organisms that act altruistically by producing public goods span the whole scale of life (Gore et al., 2009; Heinsohn & Packer, 1995; Riehl & Frederickson, 2016; Scheel & Packer, 1991; Segredo-Otero & Sanjuán, 2022; Maynard Smith, 1965; Smith & Schuster, 2019; Travisano & Velicer, 2004; Ueda et al., 2012; van Dijk et al., 2014; Wang & Lu, 2018; West et al., 2007). However, because public goods production is costly while their benefits are freely available to all members of a group, such altruism is perpetually threatened by the emergence of cheaters who benefit from free access without contributing to their production. Theory suggests that this ‘public goods dilemma’ should lead to the tragedy of the commons and the resulting collapse of public goods production (Fehr & Gachter, 2000; Frank, 1994, 1998; Hardin, 1968; Turner & Chao, 1999). The ubiquity of public goods in nature, however, clearly indicates that altruism can be evolutionarily stable despite the intrinsic logic of the ‘public goods dilemma’. It is widely accepted that kin selection can provide a solution to this problem. In this, a gene that causes individuals to altruistically produce costly public goods can be advantageous if sufficient benefits are reaped by carriers of that same gene within the group (Frank, 2010). This idea is succinctly captured by Hamilton’s rule, which states that an altruist gene is favoured when the benefits it provides to copies of itself, given by *Rb*(i.e., the benefits of altruism, *b*, weighted by the coefficient of relatedness between actors and recipients, *R*) are greater than the cost of altruism, *c*, which can be written as a simple inequality: *Rb*> *c* (Akçay & Van Cleve, 2016; Hamilton, 1963, 1964; Maynard Smith, 1980).

While Hamilton’s rule offers a concise statement of the conditions under which altruism is favoured by kin selection, its very nature (as a simple criterion given by an inequality) limits its ability to account for several key properties of public goods altruism observed in experimental and natural systems (Allen et al., 2013; Allen & Nowak, 2015). Critically, because Hamilton’s rule will either be met or not (ignoring the case where *Rb* = *c*), the only evolutionary outcomes are that altruism is favoured (an altruistic variant should reach fixation) or it is disfavoured (an altruistic variant should be removed by selection). Consequently, Hamilton’s rule does not provide conditions that could maintain variability in levels of public goods investment, including the occurrence of altruist/non-altruist polymorphism (often described in terms of cheater/altruist coexistence; Allison, 2005; Greig & Travisano, 2004; Riehl & Frederickson, 2016; Wang & Lu, 2018). Such variation is observed widely in nature (Riehl & Frederickson, 2016; Wang & Lu, 2018), with examples including: replication enzymes in phage (Turner & Chao, 1999, 2003), quorum sensing signals (Diggle et al., 2007; Pollitt Eric J. G. et al., 2014), biofilm components (Martin et al., 2020; Nadell et al., 2016), enzymatic degradation of antibiotics (Dugatkin et al., 2004) and iron scavenging siderophores (Dumas & Kümmerli, 2012; Griffin et al., 2004; Smith & Schuster, 2019) in bacteria, fruiting body formation in social microbes (Strassmann et al., 2000; Tarnita, 2017; Velicer et al., 2000), extracellular invertase in yeast (Gore et al., 2009; Greig & Travisano, 2004), ‘royal cheats’ in ant colonies (Dobata & Tsuji, 2013; Hughes & Boomsma, 2008), and contributions to group hunting in predators such as lions (Heinsohn & Packer, 1995; Scheel & Packer, 1991).

The most widely accepted explanation for cheater maintenance is that the payoffs for altruists versus cheaters match the snowdrift game (SD), where cheating (‘defection’) is the best strategy against an altruist (often described as ‘cooperating’) while altruism is the best strategy against a cheater (Doebeli & Hauert, 2005; Gore et al., 2009; Turner & Chao, 2003). A number of game theoretical models have investigated the conditions that promote or limit such diversity, with the exact outcomes depending on the existence of synergistic effects (Doebeli et al., 2004; Fletcher & Doebeli, 2006; Hauert et al., 2006; Sasaki & Okada, 2015) and on the presence of spatial structure (Hauert & Doebeli, 2004, 2021). Importantly, it has been shown that the payoff structure could itself evolve as selection shapes altruist and cheater strategies (Chao & Elena, 2017; Doebeli et al., 2004; Sasaki & Okada, 2015). Therefore, understanding the biological factors that dictate the payoff structure is critical to explain patterns in nature and outcomes in experimental studies (Doebeli & Hauert, 2005; Gore et al., 2009; Turner & Chao, 2003).

One of the key factors that should govern the nature of payoffs for altruism or cheating is how altruistic the altruists actually are — i.e., how much of their resources (e.g., time and energy) they invest into public goods (Chao & Elena, 2017; Doebeli et al., 2004). The level of resource investment into public goods should reflect the direct fitness costs and the inclusive fitness benefits (owing to relatedness), such that different investment levels would be expected to evolve when organisms face different conditions. This has been observed in viruses, bacteria, fungi, and facultatively social insects (Bastiaans et al., 2016; Griffin et al., 2004; Keller, 2003; Turner & Chao, 1999, 2003; Walter & Heinze, 2015). One limitation of existing models is that they typically treat the costs and benefits of altruism as effectively arising from two-separate phenomena. However, if there are trade-offs over the use of limited resources, then the costs and benefits should be functionally linked. When individuals invest their limited resources into public goods, they necessarily diminish the potential fitness benefits they can receive from those public goods. This is because they have sacrificed some of the resources needed to realise the benefits (e.g., by increasing survivorship) (Bachmann et al., 2013; Lee et al., 2016; Michod, 2006; Wolf et al., 2015). Consequently, the inherent resource allocation trade-offs that organisms face when investing into altruism are expected to play a critical role in shaping public goods altruism, yet are rarely considered (Roff, 1993; Stearns, 1989).

Here, we integrate the logic of resource allocation trade-offs and kin selection theory to formulate a powerful, biologically motivated framework for modelling public goods dilemma. This allows us to derive general solutions that simultaneously address the two key outstanding problems: how altruistic should individuals actually be and when can cheaters coexist with altruists. By treating public goods altruism as a quantitative trait that evolves in the face of resource allocation trade-offs, our approach provides a critical bridge to game-theoretical models, where the payoff structure for altruism and cheating is a consequence of the traits, not a property of the model itself. Key to our framework is the development of a flexible model of group-structured social interactions that can capture the broad array of phenomena that contribute to variation in group composition (and hence relatedness), including population structure (reflecting phenomena such as population viscosity), shared ancestry, partner choice, and group size (Balding, 2003; Balding & Nichols, 1995; Rannala & Hartigan, 1995). By capturing hierarchical population structure, this model also provides a straightforward means of evaluating the impact of local competition on evolutionary outcomes. Our framework provides easily interpretable results that can be directly related to properties of biological systems. Consequently, these results can be used to develop explicit testable predictions and enable researchers to develop a full understanding of public goods altruism in nature.

## Model and Results

We assume that public goods altruism is determined by a single haploid locus with two alleles: an altruist allele ‘*A*’ (with frequency *p*_*A*_), and a non-altruist allele ‘*N*’ (with frequency *p*_*N*_ = 1 − *p*_*A*_). We further assume that individuals interact in groups, where the level of public goods available in that group influences the fitness of all members of the group (i.e., benefits of public goods are freely accessible to all group members). We formalise this theoretical framework by first describing how we model the distribution of allele frequencies across interaction groups. We later generalise the model of group structured interactions to consider the case of a hierarchically structured metapopulation, which allows us to include local competition. We next outline the basic premises of our model for how public goods investment affects fitness by first considering a case of additive costs and benefits. This allows us to describe the evolutionary properties under a scenario that follows the same basic assumptions as those underlying Hamilton’s rule (Hamilton, 1963; Queller, 1985; Taylor, 1992). Thereafter, we extend this fitness model to include the resource allocation trade-off that results from individuals either investing their limited resources to produce public goods or retaining those resources to further their own direct fitness.

### A model of group-structured interactions

Following the basic logic of trait-group models (Peña et al., 2015; Wilson, 1975, 1977, 1979) we assume there is an infinitely large population (which simply means we can ignore the impact of drift on allele frequencies) in which individuals interact in groups of size *G* each generation (a set of all notations can be found in Supporting Text S0 of SI). Alleles are sampled into such groups by a combination of deterministic and random processes. The deterministic processes represent those that lead to biased allele sharing among groupmates, which can arise from any form of social partner choice/preference that increases allele-sharing among social partners (groupmates) and/or from a simple bias towards interacting with relatives (Gardner et al., 2011; Pepper, 2000; Queller, 1985). We refer to this total biased allele sharing among groupmates as ‘assortment bias’ and denote it by *R*_*GT*_ (0 ≤ *R*_*GT*_ ≤ 1). The subscript *GT* refers to **g**roups relative to the **t**otal population and is introduced here for consistency with later sections, where relatedness is partitioned analogously to classical fixation indices (Pamilo, 1984; Wright, 1949). Under this formulation, *R*_*GT*_ simply measures the extent to which groupmates share the same allele more than expected by random chance, given the allele frequency in the total population.

We model this deterministic influence on group compositions using the beta distribution (following the approach of Balding & Nichols, 1995; Rannala & Hartigan, 1995), which describes the probability density of the expected frequency of altruists in groups (*p*_*A*(*G*)_) given the global allele frequencies (*p*_*A*_ and *p*_*N*_) and overall degree of assortment bias *R*_*GT*_:

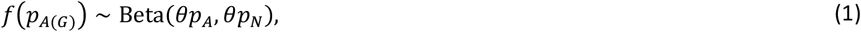

where *θ* = (1/*R*_*GT*_) − 1 (see Figure 1).

**Figure 1.**
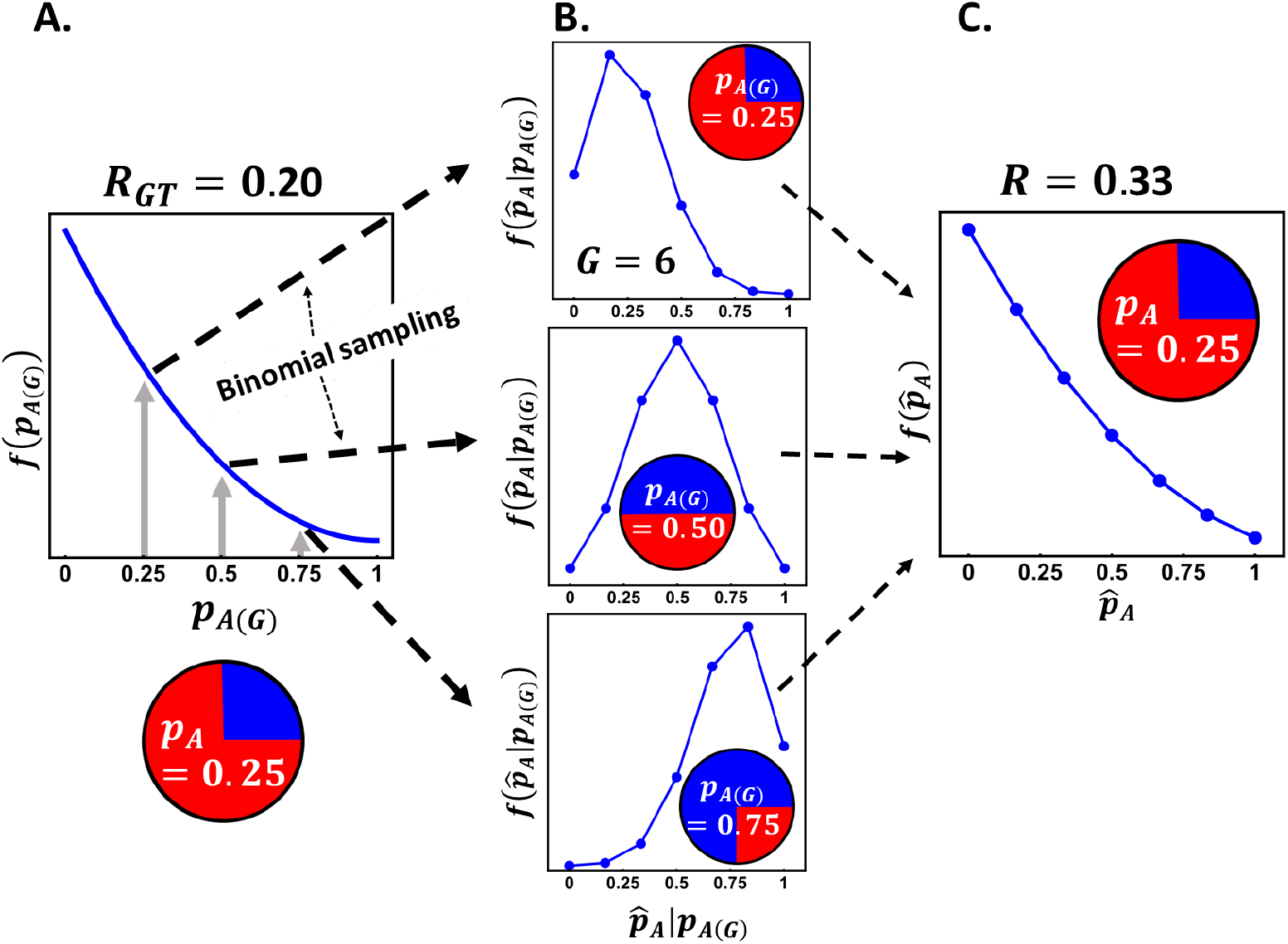
Model of group-structured interactions. Variation in the frequency of an altruist allele across groups is modelled using a beta-binomial distribution. The altruist allele has a global frequency *p*_*A*_ (illustrated here using *p*_*A*_ = 0.25. **A)** The probability density distribution of the expected frequency of the altruist allele within groups (*p*_*A*(*G*)_) is given by a beta distribution, with the shape depending on the degree of assortment bias (measured by *R*_*GT*_, illustrated here by *R*_*GT*_ = 0.20). Three expected frequencies of the altruist allele are sampled from this beta distribution, corresponding to *p*_*A*(*G*)_ = 0.25, 0.50, and 0.75. **B)** For each of these expected frequencies of altruists in a group (*p*_*A*(*G*)_), sets of alleles are binomially sampled into groups of size *G* (illustrated here with *G* = 6). Hence, each of the three plots in part B corresponds to a binomial distribution of group compositions 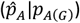, where the sampling probability is given by *p*_*A*(*G*)_ and the number of samples by *G*. **D)** These two processes (sampling of an expected allele frequency from the beta distribution and binomial sampling from that frequency to form a group of size *G*) are combined into a single beta-binomial distribution, which gives the overall probability density distribution of group compositions (in terms of the frequency of altruists in the group, 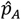), where *R* = 0.33.

If groups were infinitely large, then this expected distribution would directly yield the observed distribution of group compositions. However, the observed composition of a given finite group can deviate from this expected composition owing to the process of finite sampling. For example, if groups are full sibships, then the frequency of alleles in the parents gives the expected frequency in their offspring, but the observed frequency of alleles in the sibship depends on Mendelian sampling. We model this process by assuming that each of the *G* alleles in a group represents a Bernoulli trial with probability *p*_*A*(*G*)_, which means that the probability density of group compositions for a given expected frequency of altruists is defined by the binomial distribution:

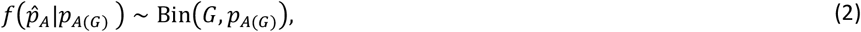

Hence, the overall distribution of compositions of finite groups will reflect both the distribution of expected group compositions (eqn. 1) and the random deviations from this distribution caused by binomial sampling (eqn. 2). These two processes can be easily combined to define the probability density function of groups with a given allele frequency 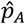 using the beta-binomial mixture distribution (Figure 1, see also Figures S1 and S2 for more details):

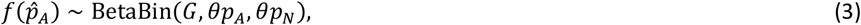

In the Supporting Text S1 we provide a more detailed description of the beta-binomial distribution and illustrations of the distribution of group compositions across different sets of parameters.

#### Whole group relatedness in a metapopulation

To link our model to the kin selection framework, we can use this distribution of group compositions to derive the relatedness, *R*, of individuals to their group (corresponding to ‘whole group relatedness’). From the perspective of the altruist allele, whole group relatedness measures the difference in the average frequency of altruist alleles experienced by altruist alleles (over all the groups) compared to that experienced by non-altruist alleles. This value can be derived as the slope from the regression of the frequency of the allele in a group 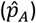 on the frequency of the allele in individuals within the group (*g*_*i*_), where the latter has a value of either 1 or 0 for the alternate *A* and *N* genotypes respectively (Michod & Hamilton, 1980; Orlove & Wood, 1978; Seger, 1981):

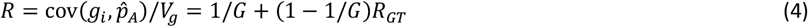

where *V*_g_ is the genetic (allelic) variance, which is equal to *p*_*A*_*p*_*N*_. Importantly, the numerator in the regression given by equation (4) represents the among-group genetic variance, which is simply *V*_*among*_ = *RV*_*g*_; hence, *R* also represents the proportion of the total allelic variation that is among groups. By extension, the within-group genetic variance is *V*_*within*_ = (1 − *R*)*V*_*g*_. The first term on the final RHS of equation (4), 1/*G*, captures the fact that, because the focal individual represents a proportion of 1/*G* of the group, their relatedness to their group will necessarily include the 1/*G* relatedness to themselves. The second term, which applies to the remaining 1 − 1/*G* proportion of alleles, represents (non-self) ‘assortative relatedness’ (Pepper, 2000) captured by the parameter *R*_*GT*_ (see equations 1 and 3).

### Evolution of public goods altruism

We assume that an altruist invests a proportion *I* of its total pool of resources into production of public goods (meaning 0 ≤ *I* ≤ 1). This comes at a personal cost to the actor of *c* (per unit invested) and provides a benefit *b* (per united invested) to the group (which gets diluted according to group size by a factor 1/*G*). The total benefit of public goods to members of a group is, therefore, the benefit provided to the group by an altruist (*bI*) times the proportion of individuals in the group that are altruists, 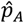, making the total benefit to all members of the group 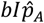. As noted above, we consider two models for how costs and benefits are combined to determine total fitness, one that assumes they are additive, and one that considers how they capture resource allocation trade-offs.

#### Evolution of public goods altruism with additive costs and benefits

Under the assumption of additivity, absolute fitness of an individual, *w*_*i*_, depends simply on the sum of the costs paid for investing in public goods (*cI* for the altruist and 0 for the non-altruist) and the benefits arising from the total level of public goods produced by all members of their group 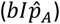. We assume for simplicity that baseline fitness is scaled to a value of 1 (which imposes a constraint that *cI* < 1 to ensure that fitness is non-negative), such that the fitness of altruists 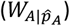 and non-altruists 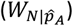 in a group containing a proportion 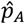 of altruists are:

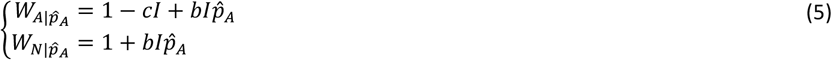

From equation (5) it is evident that the altruist allele is always disfavoured within a group because it pays the costs for the public goods that equally benefit both itself and the non-altruist allele. Therefore, for selection to favour the altruist allele, its disadvantage within groups has to be outweighed by the advantage it confers to its group (*cf*. trait group models; Wilson 1977).

To identify the conditions where the altruist allele is favoured, we derive the change in the frequency of the altruist allele. For this, we first calculate the selection coefficient for the altruist allele: 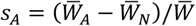, where 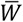 is the average fitness in the population and 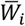 represents the mean fitness of each of the alleles over all groups: 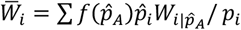 (van Veelen, 2018). The selection coefficient can also be calculated as the slope of the regression of individual relative fitness 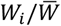 on the frequency *p*_*i*_ of the altruist allele in the individual (Gardner, 2020; McGlothlin et al., 2014). Following either approach yields a selection coefficient that has a form analogous to Hamilton’s rule: 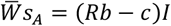, except that the cost-benefit differential (*Rb*− *c*) scales with the level of investment *I* and the relatedness term refers to whole-group relatedness, which includes self. The change in the frequency of the altruist allele can be calculated from the selection coefficient times the allelic variance, *s*_*A*_*V*_g_:

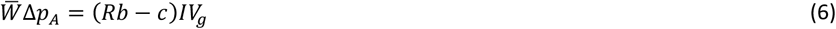

where 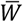 is the overall mean fitness of the population 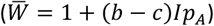. Because the selection coefficient does not depend on the frequency of the alleles, there are no polymorphic evolutionary equilibria (i.e., the altruist is either favoured or disfavoured, and hence the only evolutionary outcomes are either fixation or loss). In the Supporting Text S2 we provide an alternative derivation of equation (6) based on the Price equation (i.e., the covariance between the frequency of the altruist allele in individuals and their relative fitness: 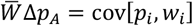.

The appearance of whole-group relatedness in equation (6) means that the benefit term (*Rb*) can be decomposed into a direct fitness component, *b*/*G*, which captures the self-benefit received by an altruist from their own altruism, and an inclusive fitness component, *bR*_*GT*_(1 − 1/*G*), which reflects the benefit they provide to other copies of the altruist allele in the (1 − 1/*G*) non-self proportion of the group. The self-beneficial effect of public goods altruism can facilitate the conditions for the invasion of an altruistic allele (by reducing the level of relatedness among group mates and/or the benefit to cost ratio required for invasion by the altruist allele), but it gets rapidly eroded/diluted as group size increases (Fletcher & Zwick, 2004; Pepper, 2000). Note that there are cases where public goods appear to rely on other-only relatedness, which our framework can easily accommodate by removing the direct fitness component. Importantly, the scaling of the selection coefficient with investment implies that, where *Rb*− *c* > 0 (i.e., Hamilton’s rule is met), the relative fitness of altruists indefinitely increases with investment, implying that relative fitness is maximised as *I* → 1 (i.e., altruists should invest all of their resources into altruism). This intrinsic shortcoming of the additive model stems from the absence of any trade-off in the definition of fitness.

#### Evolution of public goods altruism with resource trade-offs

Since public goods production requires investing resources that might otherwise be used to enhance direct fitness, we focus on a scenario where individuals have a fixed resource budget from which the investment is ‘withdrawn’ (and hence unavailable for other uses). In this scenario, the benefit of public goods comes about by increasing the expected value of the residual resource pool retained by individuals after any investment into public goods (see Madgwick et al., 2018; Queller, 1985; Maynard Smith, 1980; Uyenoyama & Feldman, 1982). This assumption follows how resource trade-offs are typically modelled in areas such as life-history theory (Michod, 2006; Roff, 1993; C. C. Smith & Fretwell, 1974; Zera & Harshman, 2001). We capture this inherent trade-off in public goods by adapting the fitness model from the Collective Investment game (Madgwick et al., 2018). The fitness values of altruists 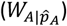 and non-altruists 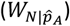 in a group containing a proportion 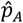 of altruists are (see also Madgwick & Wolf, 2020):

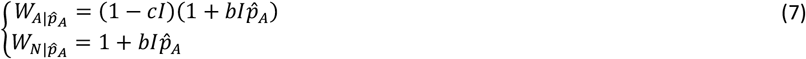

Following the approach outlined for the additive scenario above, we can derive the selection coefficient, *s*_*A*_, for the altruist allele as the difference in the average relative fitness of the two alleles over all groups and use that to calculate the change in the frequency of the altruist allele:

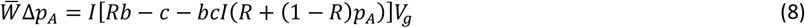

From equation (8), we can see that the same cost-benefit term appears as for the additive case (*Rb*− *c*; eqn. 6), plus an additional term, *bcI*[*R* + (1 − *R*)*p*_*A*_]. This extra term represents the opportunity cost generated by the trade-off, whereby altruists forgo some of the direct fitness benefit they could otherwise reap from public goods by diverting resources into producing them. The component *I*[*R* + (1 − *R*)*p*_*A*_] represents the average level of public goods investment experienced by the altruist allele. Importantly, this quantity increases with *p*_*A*_, introducing a negative frequency-dependent component of selection whose strength declines with relatedness. Overall, equation (8) shows that a necessary condition for altruism to invade (i.e., Δ*p*_*A*_ > 0) is that *Rb*− *c* > 0. However, unlike the additive case, this is not a sufficient condition because the opportunity cost makes the conditions favouring the altruist allele contingent on the level of public goods investment, which itself depends on *p*_*A*_.

Using equation (8) we can solve for the altruist allele frequency at which the change in the frequency of the altruist allele is zero (i.e., Δ*p*_*A*_ = 0; which also corresponds to the condition where *s*_*A*_ = 0):

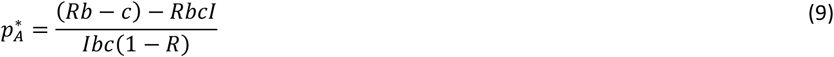

where *R* denotes whole-group relatedness. From equation (9) we can see that the altruist allele will be universally favoured over the non-altruist allele 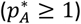 whenever *I* ≤ (*Rb*− *c*)/*bc*, and will be universally disfavoured 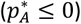 whenever *I* ≥ (*Rb*− *c*)/*Rbc*. In the parameter space between these two boundaries, (i.e., where (*Rb*− *c*)/*bc* < *I* < (*Rb*− *c*)/*Rbc*), there will be a stable polymorphic equilibrium at the allele frequency given by 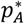 (see Figure 2). The identification of conditions favouring a monomorphic or polymorphic outcome is one of the key results of our model. Below we relate the conditions in equation (9) to the factors shaping the evolution of investment in altruist. Together, these results provide potentially important biological predictions for the scenarios where we would expect to see stable co-occurrence of altruists and cheaters in nature.

**Figure 2.**
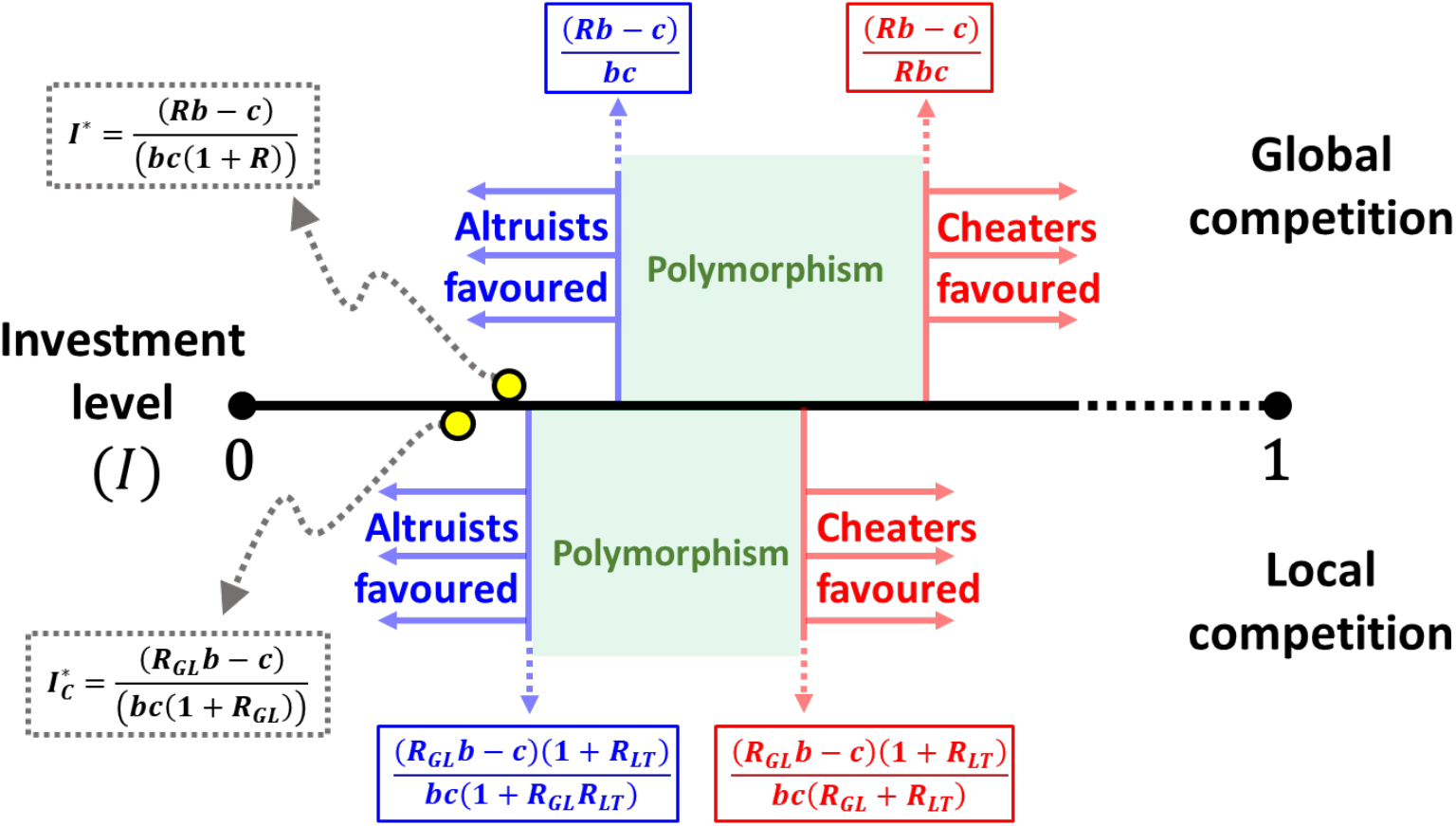
Critical investment values under global and local competition. The horizontal line represents the level of investment made by an altruist (from zero to some intermediate level, with the maximum value of 1 being further to the right on the real scale). Above the horizontal line are results for the model under global competition (i.e., no competitive constraint), while below the line are results for the scenario where there is local competition. Starting from the left, the yellow circles indicate the optimal (ESS) level of investment by an altruist (with arrows pointing to the corresponding equations), which will be higher under global competition than local competition (assuming the same values for *R*_*GL*_ and *R*_*LT*_ under the two scenarios). Next is the investment threshold (in blue) below which altruists are unconditionally favoured (over non-altruists = cheaters). Above that investment threshold is a range of investment levels where polymorphism between altruists and cheaters can be maintained (shaded region in green). That region ends at the threshold (in red) above which cheaters are unconditionally favoured so that altruism cannot invade. As can be seen in the equations, the exact positions of these thresholds will depend on the values of *c, b*, and the relatedness values (*R*_*LT*_, *R*_*GL*_ and hence *R*). The ordering of the thresholds (but not the exact distances between them) illustrated here corresponds (approximately) to a scenario where *c* = 1, *b* = 5, *R*_*GL*_ = 0.5, and *R*_*LT*_ = 0.1.

##### Within- and among-group selection on altruism

To help provide a conceptual understanding of the nature of selection on public goods altruism, and hence the conditions that lead to polymorphism, we can decompose selection on the altruist allele into components that reflect within- and among-group selection (see Supporting Text S2 for details on the derivation of these components). The within-group component gives the change in the frequency of altruists within groups:

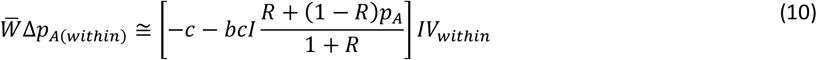

while the among-group component gives the change in the frequency of altruists that results from the differential fitness of groups caused by variation in their frequency of altruists:

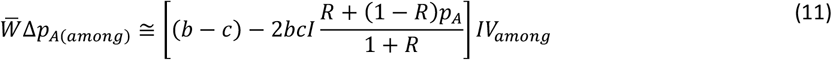

These two equations capture how costs, benefits, and the opportunity cost differentially contribute to within- and among-group selection. We see that, the cost of investing contributes negatively to both components of selection, while the benefit from investment only contribute to the among-group component. As with overall selection (eqn. 8), the opportunity cost term in both components shows negative frequency dependence, with the degree of frequency-dependence decreasing with relatedness. Importantly, the opportunity cost term in the among-group component is multiplied by 2, indicating that the opportunity cost plays a larger role in among-group selection (though the weighting of within- and among-group selection will ultimately depend on the size of *V*_*within*_ vs *V*_*among*_). Note that the exact analytical solutions (given by eqns. S8 and S9) differ from these approximations in the form of the opportunity cost term. This difference arises because of how the components of relatedness to self (1/*G*) and assortative relatedness (*R*_*GT*_) contribute to the opportunity cost. This reflects the fact that the opportunity cost arises solely from the reduction in an individual’s ability to exploit the public goods produced by others. Their own investment in public goods does not contribute to the opportunity cost because, if they were to retain those resources instead, their personal public goods contribution would disappear.

Using equations (10) and (11), we can identify the conditions where the within- and among-group components of allelic frequency change are equal and opposite, which returns the evolutionary equilibrium of eqn. (9) (see also Wade, 1985; Wade & Breden, 1980). Hence, the altruist and non-altruist alleles can both be retained in a population when the strength of selection disfavouring altruists within groups is perfectly counterbalanced by the strength of selection favouring altruists among groups.

##### Evolution of the quantity of investment into public goods

Because the conditions for the persistence of polymorphism depend critically on the level at which altruists invest (*I*), evolutionary changes in the level of investment by altruists could logically feedback on the maintenance of polymorphism. To address this problem, we evaluate how selection should shape investment in public goods. Evolutionary changes in the level of altruism requires there to be competition between altruists that invest at different levels. Therefore, we evaluate this problem by considering competition between alternative altruist alleles, *A*_1_ and *A*_2_, with frequencies *p*_1_ and *p*_2_, that invest different levels of resources in public goods: *I*_1_ = *I* + *δ* and *I*_2_ = *I*. The fitness of each of two alleles follows the form of the fitness of the altruist allele in equation (7), modified to account for the level of investment of both alleles, such that the fitness (within a group) of allele *A*_i_ is 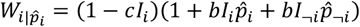 (where ¬*i* is read as ‘not *i’*, meaning it refers to the alternative allele). To evaluate whether an allele is favoured over an(y) other, we calculate the frequency change of allele *i* as 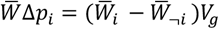, which yields 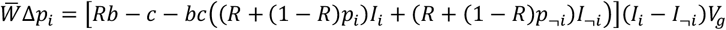 for the general case, and 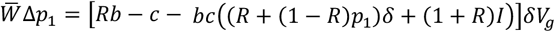 for allele *A*_1_ (where Δ*p*_2_ = −Δ*p*_1_ if it competes against *A*_2_).

Solving for the level of investment where the evolutionary change in the frequency of allele 1 is zero, denoted 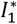, yields:

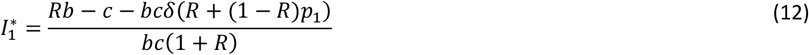

Taking the value of 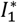 at the limit where *δ* → 0 makes investment by *A*_1_ and *A*_2_ indistinguishable, yielding the evolutionarily stable level of investment *I*^*\**^ that is uninvadable:

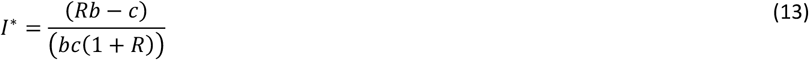

In the Supporting Text S3 we use adaptive dynamics to prove that this level of investment is an evolutionarily stable strategy (ESS) that cannot be invaded by a mutant showing any different level of investment, in agreement with findings for family-structured interactions (Uyenoyama & Feldman, 1982), and that it is also convergent stable as it can be approached through gradual mutations.

To determine the cause behind the existence of an optimal level of investment, we can rearrange equation (13) as *bc*(1 + *R*)*I*^*\**^ = *Rb*− *c*, which illustrates the fact that, at the ESS level of investment, the opportunity cost for investing in public goods, *bc*(1 + *R*)*I*^*\**^, exactly offsets the simple benefit-cost differential of Hamilton’s rule. Logically, the optimal level of investment increases with *R* and *b*, and decreases with *c*. Because the value of relatedness (*R*) in equation (13) represents whole-group relatedness (in our framework), and therefore includes the contribution of the relatedness of the individual to itself, the optimal level of investment also logically increases with smaller groups, all else being equal (since the actor gains a larger direct benefit from its own contribution to public goods).

##### Relationship between investment levels and maintenance of polymorphism

Given a particular set of values for relatedness and the costs and benefits of public goods investment, the conditions that allow for the maintenance of polymorphism (see eqn. 9) depend critically on the level of investment, while the level of investment itself is expected to evolve in response to these same factors (see eqn. 13). Therefore, to understand the conditions where we would expect to see polymorphism, it is important to relate these two processes (maintenance of polymorphism and evolution of investment levels). If the level of investment is free to evolve to its ESS (eqn. 13), then, by definition, non-altruists would be excluded. Hence, the conditions that can allow for polymorphism must require altruists to invest at a level that is not the ESS. Such a scenario could arise if evolutionary constraints limit the possible quantities of investment that individuals can make into public goods, such as might be the case if the altruistic act is an all-or-none behaviour (e.g., an alarm call, or helpers sacrificing a reproductive season).

To analyse this problem, we again consider investment by competing alleles (*A*_1_ and *A*_2_), but we now measure their investment as deviations from the optimal level of investment: *I*_1_ = *I*^*\**^ + *δ*_1_ and *I*_2_ = *I*^*\**^ + *δ*_2_. Calculating the selection coefficient on the alleles as above, the conditions where the selection coefficient for allele *A*_1_ is zero yields:

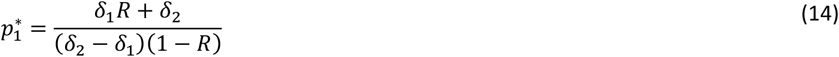

From equation (14) it can easily be shown that, if both *δ*_1_ > 0 and *δ*_2_ > 0 (i.e., both alleles invest above the optimum) or both *δ*_1_ < 0 and *δ*_2_ < 0 (i.e., both alleles invest below the optimum), then there will be a monomorphic equilibrium, where the allele closer to the optimum (i.e., with the deviation closer to zero) would go to fixation. Therefore, the only conditions that can potentially allow for the persistence of polymorphism are those where one allele is an ‘overly generous’ altruist that invests above the optimum and the other is a ‘stingy’ altruist that invests below the optimum, which logically includes the extreme case of an altruist and a non-altruist (cheater). Whichever allele is closer to the optimum will be the more common allele, regardless of whether it is the one above or below the optimum. Finally, if a third allele *A*_3_ were to arise and its level of investment was closer to the ESS than the existing allele that is on the same side of the optimum (e.g., *A*_2_), then it would replace that allele and it would reach an equilibrium frequency with the remaining allele (in this case, *A*_1_) that is necessarily higher than that of the allele it replaced since its investment is closer to the optimum (see Supporting Text S3).

### Impact of the scale of competition on public goods altruism

For the benefits of public goods altruism to be realised, they must increase the fitness of a group relative to other groups. Consequently, the presence of competitive constraints that equalise average fitness across different areas in a hierarchically structured population could logically remove some of the benefits of public goods. Analyses of the impact of local competition based on models with additive costs and benefits have shown that this can fully preclude the evolution of altruism by negating the indirect component of inclusive fitness that relies on the benefits to relatives (Queller, 1994; Taylor, 1992; West et al., 2002). To understand how such local competition impacts the evolution of public goods altruism, we next consider how the scale of competition, shifting from global to local, impacts each of the outcomes of our model.

#### Components of relatedness and the scale of competition

We can assess the impact of the scale of competition on evolutionary outcomes by modifying our model of group-structured interactions to allow for ecological constraints that differ across levels of population structure. Accordingly, we examine the scenario where groups can vary in fitness within local areas (the lowest level of population structure), yet all local areas are constrained to the same mean fitness. This assumption implies that local areas all have the same productivity, which could occur owing to competition for a fixed quantity of resources. Because we do not impose such a competitive constraint in our derivation of the model above, it implicitly assumes ‘global’ competition amongst all groups. By contrast, when a competitive constraint forces local areas to have equal mean fitness, competition is restricted to occurring only among groups within the same local area, which we refer to as ‘local’ competition (Griffin et al., 2004; Queller, 1994).

The scale of competition can be captured through the partitioning of assortment bias (*R*_*GT*_) into two components *R*_*GL*_ and *R*_*LT*_, each of which corresponds to the distribution of allelic variation at the scale at which competition occurs. We can achieve this separation in our model by replacing the beta distribution (used to model assortment bias above) with nested beta-distributions, where each accounts for the assortment bias existing at a given level. We assume that, at the highest level of population structure, allelic variation can be unevenly distributed across local areas owing to phenomena that lead to patchiness, such as population viscosity or physical barriers to gene flow. We label this source of allele frequency variation as *R*_*LT*_, where the subscripts indicate **L**ocal areas relative to the **T**otal metapopulation. This parameter will typically correspond to the standard population genetic parameter *F*_*ST*_, and can therefore be related to migration rates in island models (Balding, 2003; Wright, 1949). We also assume that local subpopulations are infinitely large such that we can model the distribution of allele frequencies across local subpopulations (*p*_*A*(*L*)_ and *p*_*N*(*L*)_) using the beta distribution. Hence, the probability density of local allele frequencies given global allele frequencies (*p*_*A*_ and *p*_*N*_) and *R*_*LT*_ is:

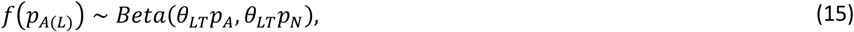

where *θ*_*LT*_ = (1/*R*_*LT*_) − 1 (and 0 ≤ *R*_*LT*_ ≤ 1). This distribution represents a metapopulation in which the variance in allele frequencies across local areas is *R*_*LT*_*p*_*A*_*p*_*N*_, while the variance within local subpopulations is (1 − *R*_*LT*_)*p*_*A*_*p*_*N*_.

We assume that, given the local allele frequency (*p*_*A*(*L*)_) that was sampled from the first distribution, interaction groups are composed by sampling alleles into groups according to the same combination of deterministic and stochastic processes described previously for the global competition scenario, except it reflects the level of assortment bias at this level, which is given by *R*_*GL*_ (where the subscripts indicate **G**roups relative to the **L**ocal area):

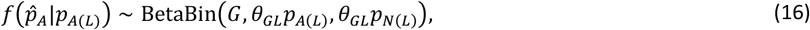

where *θ*_*GL*_ = (1/*R*_*GL*_) − 1.

The degree of assortment bias at the two levels of population structure, *R*_*LT*_ and *R*_*GL*_, together contribute to the total level of variation among groups, such that *R*_*GT*_ = *R*_*GL*_ + (1 − *R*_*GL*_)*R*_*LT*_. Hence, the overall level of relatedness would follow the form in equation (4) when the contribution of self-relatedness in finite groups is considered:

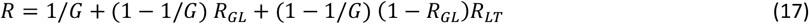

For clarity, we present the results in terms of *R*_*GL*_ and *R*_*LT*_, implicitly assuming large interaction group sizes. These results naturally extend to small group sizes if *R*_*GL*_ is replaced by the explicit form of whole-group relatedness 1/*G* + (1 − 1/*G*) *R*_*GL*_.

#### Selection in a hierarchical model of group structure

To derive the result where there is a competitive constraint, we follow the same logic described for the case of global competition, except we assume that all local areas have the same average fitness. In the absence of a competitive constraint, we could calculate the overall change in the frequency of the altruist allele by first calculating the change within local areas 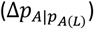 and then summing over all local areas by weighting the change within a local area by relative local mean fitness 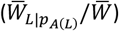. In other words, the global change in the allele frequency would be the relative fitness weighted average of the local changes. The presence of a competitive constraint effectively removes the weighting by relative local mean fitness (since all local areas have the same mean fitness), so the global change is simply the unweighted average local change: 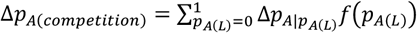. To calculate the local change in altruist allele frequency, we need to calculate the local selection coefficients, which depend on the local differences in the average relative fitness of the two alleles: 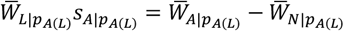 (Taylor 1992; also see Supporting Text S4). The local selection coefficient is then multiplied by the local allele frequencies to calculate the local allele frequency change, such that 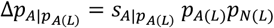. Summing over all local areas gives:

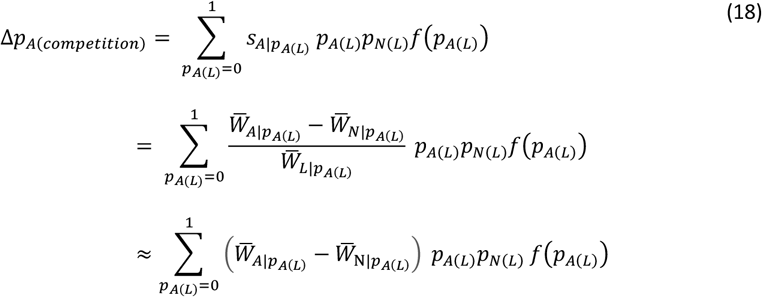

The exact solution to equation (18) involves the summation of a ratio (shown on the second line) for which there is no generic closed form solution. For simplicity and clarity, we focus on the approximation shown on the bottom line, which assumes that selection is weak enough such that we can neglect the influence of the denominator 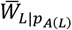. In the Supporting Text S4 we provide exact solutions and further details on the approximation shown here. Importantly, under most relevant parameter space, the expectations produced by the approximate and exact solutions are very close (see Figures S5-7).

#### Impact of the scale of competition with additive costs and benefits

To understand the consequences of local competition, we first consider the scenario with additive costs and benefits (following the fitness model given by eqn. 7). Under this scenario, the local selection coefficient is 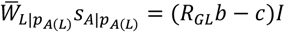, which yields a solution to equation (18) that is analogous to the case of eqn. 8) without local competition (see Supporting Text S4 for details):

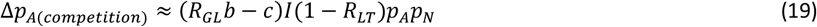

Hence, altruism is favoured whenever Hamilton’s rule is met within local areas (*R*_*GL*_*b* > *c*; (Queller, 1994). Noticeably, the component of relatedness associated with variation in allele frequencies among local areas (*R*_*LT*_) will still influence evolutionary dynamics, but only in terms of reducing the rate (by reducing the amount of genetic variation within local areas), not the direction of allele frequency change. Finally, in the absence of a competitive constraint (where *R*_*LT*_ = 0 and *R*_*GL*_ = *R*), equation (19) logically returns the expression under global competition (eqn. 6).

#### Impact of the scale of competition on the evolution of public goods altruism with resource trade-offs

We can apply the same logic as above to investigate how local competition impacts the evolution of altruism for the resource allocation trade-off scenario. Under this scenario, the local selection coefficient would be: 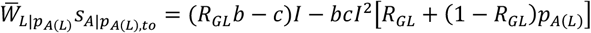. As expected from the scenario with global competition, the local opportunity cost term depends on the average level of investment experienced by the altruist allele given the local allele frequency: *I*[*R*_*GL*_ + (1 − *R*_*GL*_)*p*_*A*(*L*)_]. This term introduces frequency-dependent selection within the local area. Plugging 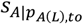 into equation (18) gives the approximate change in the frequency of the altruist allele under this scenario (see Supporting Text S4 for details):

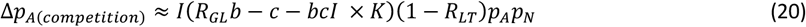

The separate impacts of benefits and costs of public goods investing match the additive case (eqn. 19). The influence of the opportunity cost term depends on 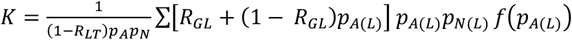. Given that local areas have large population sizes implies that *K* can be integrated over a beta distribution and is approximately equal to (see Supporting Text S4):

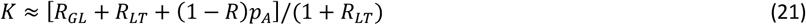

Using equation (21) in place of *K* in equation (20) we can then solve for the equilibrium frequency of the altruist allele:

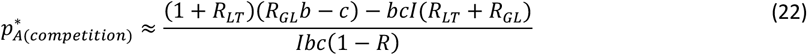

which predictably matches the equilibrium conditions in the absence of a competitive constraint when *R* = *R*_*GL*_ (and *R*_*LT*_ = 0).

Unlike the additive case, where only the group-level relatedness *R*_*GL*_ matters for the conditions favouring the altruist allele, the presence of a resource trade-off means that both components of relatedness now influence the evolutionary outcome. The altruist allele will be universally favoured (i.e., 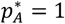) whenever *I* ≤ (*R*_*GL*_*b* − *c*)(1 + *R*_*LT*_)/[*bc*(1 + *R*_*LT*_*R*_*GL*_)], while the non-altruist allele is universally favoured whenever *I* ≥ (*R*_*GL*_*b* − *c*)(1 + *R*_*LT*_)/[*bc*(*R*_*LT*_ + *R*_*GL*_)]. Between these thresholds, there will be a polymorphic equilibrium coinciding with equation (22). These conditions have similar properties as the case with global competition but are less favourable to altruism. The threshold level of investment below which altruism is universally favoured is lower relative to the global competition case, as is the threshold level of investment above which the non-altruist is favoured. The range of investment levels between these two thresholds will also be smaller than under the global competition scenario, which means that there is less parameter space in which polymorphism can be maintained (see Figure 2).

Finally, following the same approach outlined above for the case of global competition, we can identify the ESS level of investment, which will always be lower than in the absence of local competition (see explanations in Supporting Text S3):

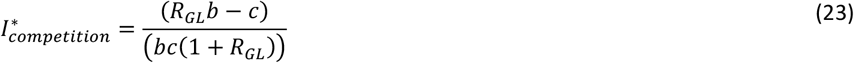

This matches the form of the ESS level of investment under global competition, except in this case only group-level relatedness matters. In both cases (local or global competition), the ESS level of investment maximises the inclusive fitness return on investment, and hence depends on the relevant measure of relatedness that determines the indirect component of fitness.

## Discussion

Kin selection is widely recognised as the key driver behind the evolution of altruism in nature. Here we extend the kin selection framework to understand the evolution of public goods altruism, where all individuals in a group benefit from the actions of altruists. For this, we developed a general model for group-structured interactions that can account for different sources of assortment bias in the composition of finite groups. We first apply this model to the ‘classic’ case of additive costs and benefits of altruism, which yields a solution for the conditions that favour altruism that matches Hamilton’s rule (*Rb*− *c* > 0), except that *R* represents ‘whole group relatedness’ (in place of the usual coefficient of relatedness; see eqn. 4), which includes the relatedness of the actor to itself (Fletcher & Zwick, 2004; Pepper, 2000). Next, we applied this approach to a more biologically grounded scenario where individuals face a strategic trade-off in the allocation of limited resources when investing into public goods production (Madgwick et al., 2018). Unlike the simple Hamilton’s rule solution, which only provides the basic conditions that favour altruism, this more realistic approach allowed us to address how much individuals (or genotypes) should be willing to invest into public goods, and to determine what conditions might allow cheaters to coexist with altruists.

When individuals invest into public goods, they necessarily reduce the amount of resource they can allocate to other components of fitness (e.g., they would have fewer resources they can use to make eggs). The presence of public goods, in return, raises the value of these residual resources (in terms of how they translate into realised fitness; e.g., they may increase survivorship of offspring), which can more than offset the reduction in the residual resource budget (Madgwick et al., 2018). As a result, this allocation trade-off entails a two-fold investment cost: altruists initially pay for the production of public goods, and by doing so, also suffer an opportunity cost by forgoing some of the potential benefits from the public goods. The presence of an allocation trade-off gives rise to optimising selection on public good investment, wherein fitness is maximised at an intermediate investment level (see eqn. 13) that represents an ESS (Uyenoyama & Feldman, 1982). Unsurprisingly, this result predicts that the level of resource investment into public goods should be higher as the relatedness and benefits increase and/or the costs decrease, which could be tested experimentally by varying these properties (e.g., in fungi, Bastiaans et al. 2016). Notably, this finding also predicts that the level of investment should be higher in smaller groups, where whole group relatedness is elevated because it includes the relatedness of an actor to itself. This phenomenon is sometimes described as ‘weak altruism’ because the actor directly benefits from the public goods they produce. Because this effect quickly erodes as group size increases (Fletcher & Doebeli, 2009), its relevance is logically restricted to very small groups.

An implied prediction of our analysis of the ESS level of investment into public goods is that, if individuals invest at this ESS level, this would (by definition) preclude the possibility of altruist/cheater polymorphism. Hence, when polymorphism is observed, we would expect to find that cheaters can persist because the altruists are being overly generous (i.e., investing above the ESS level, see Figure 2). This result is conceptually analogous to the finding of a mixed ESS in the simple all-or-none snowdrift scenario (Gore et al., 2009; Hauert & Doebeli, 2004; Sasaki & Okada, 2015; see Supporting Text S5). Such conditions could arise in nature when altruistic acts are evolutionarily constrained by being all-or-none (e.g., alarm calls, (Charnov & Krebs, 1975; Hoogland, 1983; Maynard Smith, 1965); programmed cell death in unicellular organisms, (Durand et al., 2011), or where costs come in discrete units, which prevents optimal investment (e.g., helpers in cooperative breeders sacrifice entire breeding seasons, (Riehl, 2013; Wang & Lu, 2018). Altruistic acts can also be experimentally constrained to be all-or-none, such as in competition experiments between cheater and altruist strains (Gore et al., 2009; Sanchez & Gore, 2013; Turner & Chao, 1999, 2003). Experimental conditions can also differ from those under which the level of investment in public goods would have evolved (often by design) such that they make the altruists overly generous. Of course, the possibility of a polymorphic outcome logically extends to other variants beyond ‘pure’ altruists and cheaters, such as polymorphism involving more and less generous altruists. A clear example of this comes from the experimental evolution of MS2 phages, where a full cheater that makes no contribution to altruistic production was replaced by the emergence of partial cheater that contributes a low level of altruistic production, which came into polymorphic equilibrium with the full altruist (Meir et al., 2020).

To generate and test model predictions of evolutionary outcomes, it is essential to bear in mind that public goods altruism can only be favoured by selection where public goods increase the overall success of a group relative to other groups. As has been found previously for the snowdrift game (Hauert & Doebeli, 2004) or kin selection (Taylor, 1992; West et al., 2002), our model predicts that the optimal level of investment into public goods depends solely on local relatedness, and hence can be lower than expected under localised competition (see eqn. 23 and Figure 2). More generally, this phenomenon tilts the balance of selection in favour of cheaters, which provides two key testable predictions. Firstly, under conditions that would otherwise favour polymorphism in the absence of local competition, it could lead to cheater fixation, and secondly, under conditions that would favour altruists, it could lead to polymorphism (or, under more extreme conditions, cheater fixation). A clear example of this comes from experimental modulation of both the level of relatedness and the scale of competition in the siderophore producing bacterium *Pseudomonas aeruginosa* (Griffin et al., 2004): in these experiments, high relatedness with global competition and low relatedness with local competition led to altruist and cheater monomorphism respectively, whereas high relatedness with local competition or low relatedness with global competition maintained altruist-cheater polymorphism. Consistent with the predictions discussed above, where experimental conditions may result in altruists being overly generous (compared to the level that would be favoured under those conditions), experiments that allowed siderophore production to evolve under low relatedness found that altruists evolved reduced investment in siderophore production, which made them less exploitable by cheaters (Kümmerli et al., 2015). Similar results have been found in phage coinfection experiments (Turner & Chao, 2003), and in fungi (Bastiaans et al. 2016).

The equilibrium outcomes that we derive provide a means of understanding patterns observed in natural and experimental systems, while the dynamical equations, selection coefficients, and flexible model of hierarchical population structure can be used to understand how patterns change across experimental conditions, including evolutionary changes in levels of altruism itself. Although extending our framework to fit specific systems will necessarily modify the precise results generated by the model, such changes would not be expected to alter the fundamental predictions of our model. Therefore, we believe that it provides a compelling and tractable description of a wide array of altruistic scenarios in nature, as well as a powerful tool for predicting and interpreting experimental results. Finally, our framework formalises key connections between findings from models based on kin selection, multi-level selection, and evolutionary game theory.

## Acknowledgements

This work was supported by grant funding from the Natural Environment Research Council (NE/V012002/1).

