## Supplementary information for "The evolution of public goods altruism"

### Supporting Information for 'The evolution of Public goods altruism'

#### Supporting Text S0: Set of notations

**Table S0. Set of notations used in the main document.**

| Definition | Parameter and relations |
|---|---|
| Frequency of altruists/non-altruists in the population | $p_A/\, p_N$ |
| Frequency of altruists/non-altruists in local areas | $p_{A(L)}/\, p_{N(L)}$ |
| Expected frequency of altruists/non-altruists in interaction groups | $p_{A(G)}/\, p_{N(G)}$ |
| Realised frequency of altruists/non-altruists in interaction groups | $\hat{p}_A/\hat{p}_N$ |
| Group size | $G$ |
| Genetic assortment bias of interaction groups (**G**) relative to local areas (**L**) | $R_{GL}$ |
| Genetic assortment bias of local areas (**L**) relative to the total population (**T**) | $R_{LT}$ |
| Overall degree of assortment bias | $R_{GT} = R_{GL} + (1 - R_{GL})R_{LT}$ |
| Whole group relatedness | $R = \frac{1}{G} + \left(1 - \frac{1}{G}\right) R_{GT}$ |
| Benefit | $b$ |
| Cost | $c$ |
| Level of investment of altruist allele *i* ('A$_i$') | $I_i \in [0,1]$ |
| Fitness of altruist/non-altruist alleles within groups | $W_{A\|\hat{p}_A}/W_{N\|\hat{p}_N}$ |
| Mean fitness of altruist/non-altruist alleles within local areas | $\bar{W}_{A\|p_{A(L)}}/\bar{W}_{\mathrm{N}\|p_{A(L)}}$ |
| Mean fitness in a local area with frequency $p_{A(L)}$ of altruists | $\bar{W}_{L\|p_{A(L)}}$ |
| Mean fitness of altruists/non altruists in the population | $\bar{W}_A/\bar{W}_N$ |
| Mean fitness in the population | $\bar{W}$ |
| Overall selection coefficient for allele *I* (against $\neg I$) | $s_i = \frac{\bar{W}_i - \bar{W}_{\neg i}}{\bar{W}}$ |
| Selection coefficient of allele *i* with frequency $p_{i(L)}$ in a local area | $s_{i\|p_{i(L)}}$ |
| Evolutionarily stable investment level | $I^*$ |
| Frequency of altruist at evolutionary equilibrium | $p_A^*$ |

#### Supporting Text S1: Beta-binomial distribution and population structure

In the main text we outline our hierarchical model of population structure based on the beta-binomial distribution. Here we provide further discussion of the logic and properties of the distribution.

The beta-binomial distribution is a compound distribution, where $G$ trials are sampled from a binomial distribution for which the probability of a given outcome is itself drawn from the beta distribution. The binomial sampling process captures the influence of finite random sampling into groups (akin to the Wright-Fisher model), while the beta distribution captures sources of autocorrelation amongst the alleles sampled into the same group. The use of the beta distribution to describe the distribution of genetic variation traces back to at least (Wright, 1931), and, for the specific case of a solution to the distribution of allele frequencies at drift migration equilibrium across subgroups under the island model, to (Wright, 1943). Our implementation of the beta-binomial distribution follows the specific logic described in the Balding-Nichols model for finite groups (Balding, 2003; Balding & Nichols, 1995; Rannala & Hartigan, 1995; Rannala, 1996), which provides a hypothetical distribution of allele frequency variation across (sub)populations for analyses of population differentiation. In such a metapopulation model, that variability across (sub)populations would typically be measured by the parameter $F_{ST}$. Yet, since our analysis is focused on the distribution of interacting groups, we use the parameter, $R_{GT}$, which measures relatedness of individuals within such groups that is caused by assortment bias. In this setting, the distribution of the frequency $\hat{p}_A$ of a given allele '*A*' in interaction groups is given by:

$$f(\hat{p}_A|\theta, G, p_A) = \frac{Beta(\hat{p}_A + \theta p_A, G - \hat{p}_A + \theta(1 - p_A))}{Beta(\theta p_A, \theta(1 - p_A))}, \quad \text{(S1)}$$

where $\theta = 1/R_{GT} - 1$. In this expression, $Beta(\alpha, \beta)$ denotes the Beta function, $G$ the group size, $R_{GT}$ the genetic assortment bias, and $p_A$ the frequency of allele '*A*' in the meta-population.

The shape of the beta-binomial distribution depends on these three parameters. First, group size determines the relative importance of binomial sampling (see Fig. S1). Therefore, as group size increases the importance of binomial sampling on the distribution of genetic variation in group composition declines. Consequently, the beta-binomial distribution converges to the beta distribution once group size approaches ~20 (*cf*. Fig. S1C and Fig. S1D).

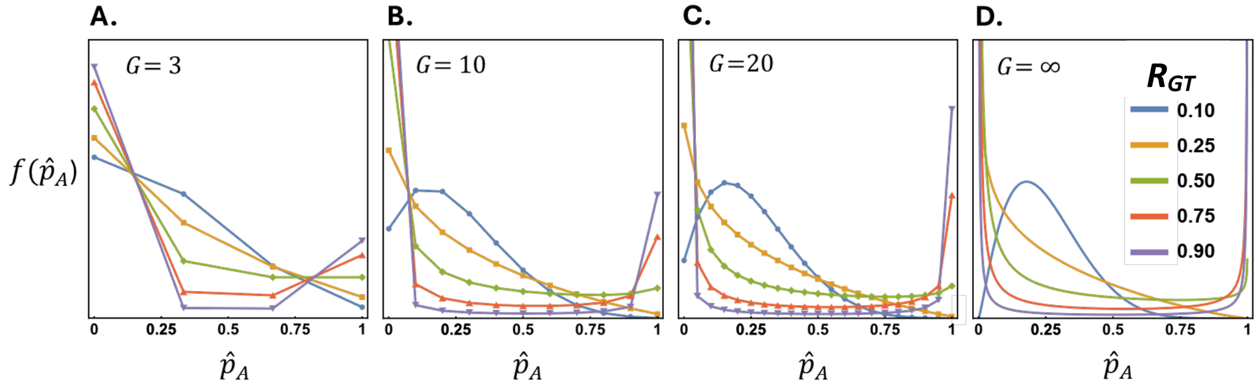


**Figure S1. Influence of group size and assortment bias $R_{GT}$ on the distribution of group composition** (in terms of the frequency of the altruist allele within groups). Each plot shows the probability density, $f(\hat{p}_A)$, of groups containing a $\hat{p}_A$ frequency of the altruist allele for different group sizes across a range of values of assortment bias. For this illustration, the overall frequency of the focal allele ($p_A$) was fixed at a value of $p_A = 0.25$. The four plots show different group sizes, corresponding to **A**) $G$ = 3, **B**) $G$ = 10, **C**) $G$ = 20 and **D**) infinite group size (which corresponds to the beta distribution). Note the similarity between $G = 20$ and the $G = \infty$, showing that large group sizes can be approximated by the beta distribution. Because the two allele frequencies sum to 1, the pattern for the non-altruist allele would simply be the mirror image of each distribution.

To understand the joint influence of the degree of biased assortment ($R_{GT}$) and allele frequencies, we illustrate a range of conditions in Figure S2 for the case of infinitely large groups (so that the influence of binomial sampling is removed).

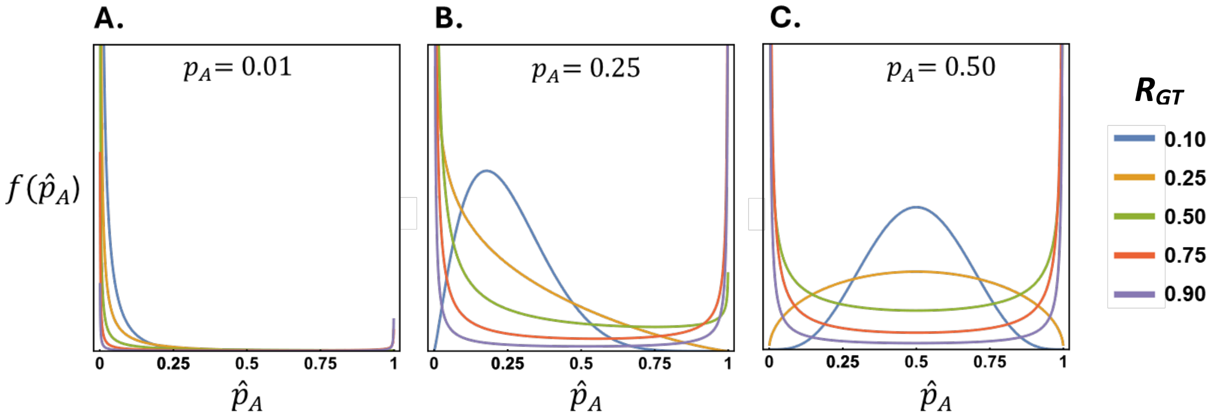


**Figure S2. Influence of the assortment bias $R_{GT}$ and allele frequencies on the distribution of group composition** (in terms of the frequency of the altruist allele within groups). Each plot shows the probability density, $f(\hat{p}_A)$, of groups containing a $\hat{p}_A$ frequency of the altruist allele for different overall allele frequencies ($p_A$) across a range of values of assortment bias. To remove the influence of group size, plots were generated using the beta distribution (implying $G = \infty$). **A**) when the altruist allele is very rare overall ($p_A = 0.01$), it tends to be very rare (no or few copies) in groups regardless of the value of $R_{GT}$. **B**) when the altruist allele is at an overall frequency of $p_A = 0.25$, the distribution is unimodal when assortment bias is very low (where $R_{GT}$ = 0.10), with probability density being maximal around the global frequency, but as assortment bias increases groups become increasingly homogeneous, with peaks at frequencies of 0 and 1. **C**) when the two alleles are at equal frequencies overall ($p_A = 0.5$), the distribution is similar to the $p_A = 0.25$ case (panel B), except the distribution is symmetrical. Because the two allele frequencies sum to 1, the pattern for the non-altruist allele would simply be the mirror image of each distribution.

To understand how selection acts on an allele, it can be useful to visualise the distribution of a focal allele across groups as the weighted probability density for an allele, where the probability of sampling a given group type is weighted by the frequency of the allele in the group, defined as $\hat{p}_A f(\hat{p}_A)/p_A$ (van Veelen, 2018). This distribution is useful because it indicates the relative importance of different group compositions for determining the overall fitness of an allele (since it illustrates the relative probability that a given allele will experience each type of group). This weighted distribution is shown in Figure S3, which provides a very different picture compared to the probability density of the different group types (*cf*. Fig. S2). For example, compare the same overall conditions illustrated in Figure S2.A and Figure S3.A. Figure S2.A shows that most groups either do not contain the focal allele, or it is at a low frequency where it occurs. However, by weighting the probability density of the groups by the frequency of the allele within groups (and re-normalising the probability so that the resulting distribution sums to 1), we see from Figure S3.A that, despite the fact that the allele is absent or rare in most groups, the average group composition experienced by copies of that allele varies widely depending on the level of assortment bias, and where $R_{GT} > 1/2$ , alleles are most likely to experience groups in which they are the most common allele.

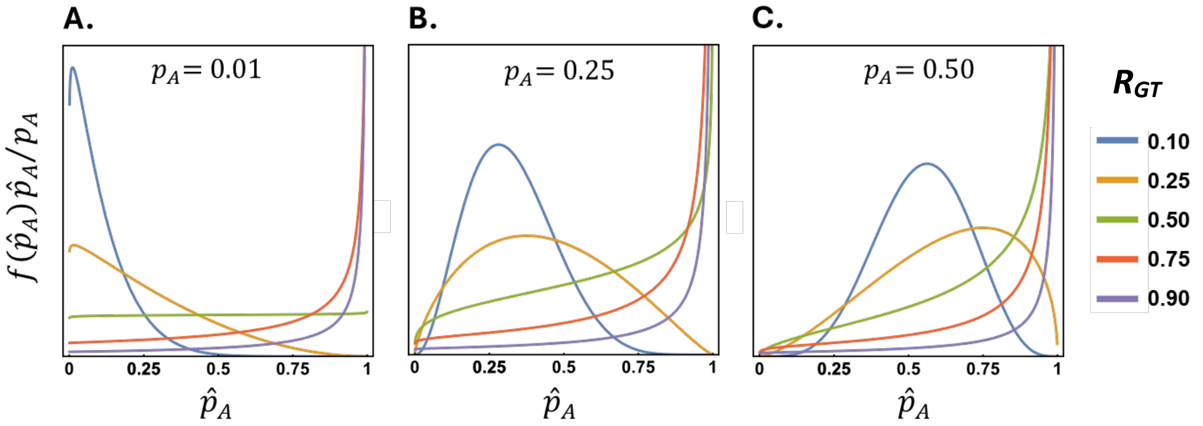


**Figure S3. Influence of the assortment bias $R_{GT}$ and allele frequencies on the weighted distribution of a focal allele across group compositions** (in terms of the frequency of the altruist allele within the group). To remove the influence of group size plots were generated using the beta distribution. The probability density, $f(\hat{p}_A)\,\hat{p}_A/p_A$, represents the relative probability that copies of the focal allele will experience each of the group compositions. **A**) when the altruist allele is very rare in the overall population ($p_A = 0.01$) and there is relatively little assortment bias ($R_{GT}$ = 0.1), most copies of the allele will be in groups where the allele itself is rare. However, when there is an intermediate level of assortment bias ($R_{GT}$ = 0.25 or 0.50) copies of the altruist allele experience a broad range of group compositions, and finally, when there is a high level of assortment bias ($R_{GT}$ = 0.75 or 0.90), most copies of the focal allele will be found in groups where the allele itself is very common. **B**) when the altruist allele is at an overall frequency of $p_A = 0.25$ and there is relatively low assortment bias ($R_{GT}$ = 0.1 or 0.25), the allele tends to experience a broad range of groups, with a bias towards its frequency in the population. However, when there is intermediate to high levels of assortment bias ($R_{GT}$ = 0.5, 0.75 or 0.90), most copies of the allele will again experience groups in which the allele is very common. **C**) when the two alleles are at the same frequency ($p_A = 0.5$), most copies of the allele will experience groups in which it is very common, even for relatively low assortment bias. The patterns for the non-altruist allele would simply be the mirror image of each of these distributions. NOTE: the y-axis in the plots in this figure have been re-scaled compared to Figure S1 and S2 to capture the patterns of variation.

#### Supporting Text S2: Using the Price equation to derive changes in allele frequencies

The evolutionary dynamics captured in the equations in the main text can also be derived using the Price equation, where the change in the frequency of an allele depends on the covariance between the frequency of the altruist allele in individuals (denoted $g_i$, where $g_i = 1$ for individuals with the altruist allele and $g_i = 0$ for those with the non-altruist allele) and their relative fitness (Gardner, 2020). Note that in sections S2 and S3, we focus on the competition between an altruist and a non-altruist allele. We then examine more complex scenarios in section S4 using Adaptive Dynamics.

##### Additive costs and benefits

For the scenario with additive costs and benefits, the Price equation covariance that gives the overall change in the frequency of the altruist allele is given by:

$$\begin{aligned}\bar{W}\Delta p_A &= \operatorname{cov}(g_i, W_i) \\ &= \sum_{\hat{p}_A=0}^{1} \left[(1-p_A)\left(W_{A|\hat{p}_A} - \bar{W}\right)\hat{p}_A + (0-p_A)\left(W_{N|\hat{p}_A} - \bar{W}\right)\hat{p}_N\right] f(\hat{p}_A) \\ &= (Rb-c)IV_g\end{aligned} \qquad \text{(S2)}$$

which matches the solution given by equation (6).

The Price equation covariance can be partitioned into within- and among-group components. The within-group covariance gives the total change in the frequency of the altruist allele within groups (so it is the weighted average across all groups). It can be calculated as the covariance between the deviation of individual fitness from the average fitness of their group (i.e., $W_{i|\hat{p}_A} - \bar{W}_{G|\hat{p}_A}$) and the deviation of the altruist allele frequency in the individuals from the frequency of the allele in their group (Wade, 1985):

$$\begin{aligned}\bar{W}\Delta p_{A(within)} &= \operatorname{cov}\left(g_i - \hat{p}_A, W_{i|\hat{p}_A} - \bar{W}_{G|\hat{p}_A}\right) \\ &= \sum_{\hat{p}_A=0}^{1} \Big[(1-\hat{p}_A)\left(W_{A|\hat{p}_A} - \bar{W}_{G|\hat{p}_A}\right)\hat{p}_A \\ &\qquad + (0-\hat{p}_A)\left(W_{N|\hat{p}_A} - \bar{W}_{G|\hat{p}_A}\right)\hat{p}_N\Big] f(\hat{p}_A)\end{aligned} \tag{S3}$$

This can be rearranged as:

$$\begin{aligned}\bar{W}\Delta p_{A(within)} &= \sum_{\hat{p}_A=0}^{1} \left(\left(W_{A|\hat{p}_A} - \bar{W}_{G|\hat{p}_A}\right) - \left(W_{N|\hat{p}_A} - \bar{W}_{G|\hat{p}_A}\right)\right)\, \hat{p}_A(1-\hat{p}_A)\, f(\hat{p}_A) \\ &= \sum_{\hat{p}_A=0}^{1} \left(W_{A|\hat{p}_A} - W_{N|\hat{p}_A}\right) \hat{p}_A(1-\hat{p}_A)\, f(\hat{p}_A)\end{aligned} \tag{S4}$$

The bottom expression captures the fact that the within-group component is the product of two terms that have simple interpretations. The first term ($W_{A|\hat{p}_A} - W_{N|\hat{p}_A}$) is the difference in the fitness of the two alleles in a group with a $\hat{p}_A$ frequency of altruists, which represents the within-group selection coefficient for the altruist allele. The value of this selection coefficient is simply $-cI$, and hence within-group selection coefficient does not depend on the frequency of altruists. The second term, $\hat{p}_A(1-\hat{p}_A)$, which is the product of the frequency of the two alleles within the group, represents the within-group allelic (genetic) variance. Logically, the product of these two terms gives the change in the frequency of the altruist allele *within* a group that has a $\hat{p}_A$ frequency of altruists. Weighting this expression by the frequency of those groups ($f(\hat{p}_A)$) and summing over all groups, therefore, gives the average within-group change in the altruist allele frequency:

$$\begin{aligned}\bar{W}\Delta p_{A(within)} &= -cI \sum_{\hat{p}_A=0}^{1} \hat{p}_A(1-\hat{p}_A)\, f(\hat{p}_A) \\ &= -cIV_w\end{aligned} \tag{S5}$$

which is the analogue version of equation (10) in the absence of resource trade-offs.

The among-group covariance gives the change in the frequency of the altruist allele across groups. It can either be calculated indirectly as the total covariance (given by eqn. 6) minus the within-group covariance of equation (S5), or directly as the covariance between the frequency of the altruist allele in a group with frequency $\hat{p}_A$ and the group's average fitness:

$$\begin{aligned}\bar{W}\Delta p_{A(among)} &= \operatorname{cov}\left(\hat{p}_A, \bar{W}_{G|\hat{p}_A}\right) \\ &= \sum_{\hat{p}_A=0}^{1} (\hat{p}_A - p_A)\left(\bar{W}_{G|\hat{p}_A} - \bar{W}\right) f(\hat{p}_A) \\ &= (b-c)I \sum_{\hat{p}_A=0}^{1} (\hat{p}_A - p_A)^2\, f(\hat{p}_A)\end{aligned} \tag{S6}$$

$$= (b - c)IV_a$$

which is the analogue version of equation (11) in the absence of resource trade-offs.

#### Resource trade-off case

The same approach as above can be used to derive the within- and among-group components of the evolutionary change in the resource trade-off case. The results for this scenario differ from the additive case because of how the opportunity cost contributes to each component. For example, the within-group selection coefficient for the altruist allele $\left(W_{A|\hat{p}_A} - W_{N|\hat{p}_A}\right) = -cI(1 + bI\hat{p}_A)$, contains both an additive cost $(-cI)$ and the opportunity cost, $-cbI^2\hat{p}_A$ terms. These two terms can be separated, such that we can write equation (S4) for this scenario as:

$$\bar{W}\Delta p_{A(within)} = -cI\sum_{\hat{p}_A=0}^{1}\hat{p}_A(1-\hat{p}_A)\,f(\hat{p}_A) - bcI^2\sum_{\hat{p}_A=0}^{1}\hat{p}_A^2(1-\hat{p}_A)\,f(\hat{p}_A) \tag{S7}$$

This separation emphasises that the opportunity cost (second term on the RHS) is weighted by a sum that differs from the additive cost in that it is multiplied by the frequency of the allele in the group. This effect arises because altruists both invest in public goods and suffer the opportunity cost from that investment (so they suffer an opportunity cost in proportion to the level of investment, which depends on the frequency of altruists in the group). This yields:

$$\bar{W}\Delta p_{A(within)} = \left[-c - bcI\frac{R + (1-R)p_A - (R - R_{GT})p_A}{1 + R_{GT}}\right]IV_w \tag{S8}$$

$$\cong \left[-c - bcI\frac{R + (1-R)p_A}{1+R}\right]IV_w$$

where the approximation on the second line is only approximate with respect to the opportunity cost term. The approximation gets more precise as group size increases, reflecting the fact that all relatedness is then due to assortment bias in infinitely large groups. Like the overall opportunity cost, the opportunity cost component of within-group selection increases with the global frequency of the altruist allele, but the influence of this frequency dependence decreases with relatedness. Neglecting the variance term, the difference between the approximation and the exact solution can be written as $(R - R_{GT})R(2p_A - 1) \times (-bcI^2)$: hence, it underestimates the size of the opportunity cost when the altruist allele is the rarer allele (i.e., $p_A < 0.5$) and overestimates the size of the opportunity cost when the altruist allele is the more common allele. Because of the $(R - R_{GT}) = 1/G(1 - R_{GT})$ term, the approximation performs better for high values of the assortment bias $R_{GT}$, and as already mentioned, for large group sizes $G$.

Following the derivation in (S6) yields the among group covariance, which is similar in structure to that of the within-group component (S8), with the major difference being that the opportunity cost is multiplied by two, indicating that the opportunity cost contributes a larger component to among-group selection than it does to within-group selection:

$$\bar{W}\Delta p_{A(among)} = \left[(b-c) - 2bcI\frac{R + (1-R)p_A - (R - R_{GT})/2}{1 + R_{GT}}\right]IV_a \tag{S9}$$

$$\cong \left[(b-c) - 2bcI\frac{R+(1-R)p_A}{1+R}\right] IV_a$$

As with the within-group covariance, the approximation on the second line is only approximated for the opportunity cost term, and also gets more precise as group size increases. Like overall and within-group selection, the opportunity cost component of among-group selection increases as the global frequency of the altruist allele increases and declines with relatedness. Neglecting the variance term, the difference between the approximation and the exact solution can be written as $(R - R_{GT})(1 - 2p_A)(1 - R)/2 \times (-bcI^2)$: contrary to the within-group component, the approximation therefore overestimates the size of the opportunity cost when the altruist allele is the rarer allele (i.e., $p_A < 0.5$) and underestimates the size of the opportunity cost when the altruist allele is the more common allele. As in the previous case, the approximation also performs better for high values of $R_{GT}$ and $G$. Note finally that the difference between the exact values and the approximations for both the within- and among-group components of selection (eqns. S8 and S9) is generally very small because they only differ with respect to the opportunity cost term.

As expected, summing equations (S8) and (S9) returns the total evolutionary change regardless of group effects (see also eqn. 8):

$$\overline{W}\Delta p_{A(total)} = [(Rb - c) - bcI(R + (1-R)p_A)]IV_g \qquad \text{(S10)}$$

#### Supporting Text S3: Adaptive dynamics and the evolution of investment in public goods

To complement and further support the evolutionary analysis presented in the main text we evaluate whether the strategies are evolutionary stable and convergent using the framework of Adaptive Dynamics (AD), which contrasts the fitness of rare mutants with that of one or possibly several resident strategies (Brännström, Johansson, & Von Festenberg, 2013; Hofbauer & Sigmund, 1990; Nowak, 1990).

##### Global competition scenario

Following the analysis of competing altruist alleles in the main text, we first consider the scenario of global competition between two altruist alleles, '$A_1$' and '$A_2$', which respectively invest $I_1$ and $I_2$ of their resources in public goods. We assume that the resident allele, $A_1$, has a frequency $p_{A_1}$~1 (meaning the frequency of allele $A_2$ ~0), such that its fitness matches that of the altruist allele given by equation (5), with its investment level being $I_1$ and its expected frequency in all groups equal to 1 (i.e., substituting $\hat{p}_A = 1$ in eqn. 7). By contrast, although the frequency of the $A_2$ allele is approximately zero, its bearers will see a distribution of their own frequency across groups given by the level of whole-group relatedness (cf. Fig. S3A), $R$, such that the average absolute fitness $\overline{W}_{A_i|p_1\to 1}$ of each allele ($A_i$) in a population of resident $A_1$ is:

$$\begin{cases} \overline{W}_{A_1|p_1\to 1}(I_1) = (1 - cI_1)(1 + bI_1) \\ \overline{W}_{A_2|p_1\to 1}(I_2; I_1) = (1 - cI_2)(1 + bRI_2 + b(1-R)I_1) \end{cases} \qquad \text{(S11)}$$

with $I_1$ and $I_2$ denoting the respective levels of investment of alleles '$A_1$' and '$A_2$'. An ESS is defined by the fact that no strategy can invade a population where it is the resident strategy (Maynard Smith & Price, 1973). It is also a fitness maximum on the adaptive landscape, such that $\overline{W}_{A_1|p_1\to 1}\ (I_1 = I^*) > \overline{W}_{A_2|p_1\to 1}\ (I_2;\ I_1 = I^*)$ for all possible values of $I_2$, and thus for values of $I_2$ close to the ESS. Dropping the index $p_1 \to 1$ for the sake of legibility, we can write the invasion fitness of a mutant (close to the resident) as a second-order approximation (Diekmann, 2002) as:

$$\overline{W}_{A_2}\ (I_2; I_1) = \overline{W}_{A_1}(I_1) + (I_2 - I_1)\frac{\partial \overline{W}_{A_2}}{\partial I_2} + (I_2 - I_1)^2\frac{\partial^2 \overline{W}_{A_2}}{\partial I_2^2} \tag{S12}$$

This expression means that the fitness of a mutant with a slightly different investment than the resident is given as a function of this slight difference and the derivatives of the invasion fitness with respect to the investment of the mutant. No mutant should be able to outperform the ESS, which means that $\overline{W}_{A_2}$ has to reach a maximum of $\overline{W}_{A_1}(I_1 = I^*)$ when the mutant $A_2$ coincides with the ESS ($I^*$). This, in turn, sets the conditions that the ESS has to fulfill: the first order derivative has to be zero (otherwise, the fitness of a mutant would be higher by investing either more or less than the resident), and the second derivative has to be negative so that any change in investment from the ESS results in being outperformed by this latter.

The first condition gives singular strategies $I^*$ that correspond to the investment level for which the invasion fitness is constant in the vicinity of the resident, that is when $I_2 = I_1 = I^*$:

$$\frac{\partial \overline{W}_{A_2}}{\partial I_2}|_{I_2=I_1=I^*} = 0 \ \Rightarrow\ I^* = \frac{Rb - c}{(1+R)bc} \tag{S13}$$

The singular strategy is an ESS if invasion fitness reaches a local maximum in this point. For this condition to be met, the resident has to satisfy the uninvadability criteria of an ESS, i.e. it has to outperform any mutant. This corresponds to the second condition defined above and is verified in our case:

$$\frac{\partial^2 \overline{W}_{A_2}}{\partial I_2^2}|_{I_2=I_1=I^*} = -2bcR \tag{S14}$$

The last condition for a strategy to evolve is that it is convergent, which means that such a (CSS) strategy can invade any other (resident) strategy (Diekmann, 2002; Eshel, Motro, & Sansone, 1997). Mathematically, this implies that $\overline{W}_{A_2} > \overline{W}_{A_1}$ when the mutant strategy is the ESS, that is $I_2 = I^*$, and the resident strategy is all but the ESS. Plugging values of equation (S11) in the fitness difference for a mutant competing against a resident yields:

$$\Delta\overline{W}_{2-1} = \overline{W}_{A_2} - \overline{W}_{A_1} = (I_2 - I_1)\big(Rb - c - bc(RI_2 + I_1)\big) \tag{S15}$$

where the mutant is considered to play the ESS, so that $I_2 = I^*$. By substituting this latter condition (eqn. S13) in equation (S15) it is then trivial to simplify the expression as:

$$\Delta\overline{W}_{2-1} = bc(I^* - I_1)^2 > 0 \tag{S16}$$

which demonstrates that the ESS is also convergent (it can invade all other strategies; see Fig. S4A). The expected adaptive outcomes are summarised in Figure S4 through the analysis of pairwise invasibility plots. In line with the analysis where competition occurs only between an altruist allele and a non-altruist one, we see that there exists a region of parameter space where a rare mutant can invade without going to fixation (yellow area in Fig. S4B). Such an area where mutual invasion is possible implies the establishment of a polymorphism, either transiently or permanently. The width of the mutual invasion area, which enables polymorphism, critically depends on $R$, vanishing when $R \to 1$ or predominating when $R$ is very low (but where $Rb - c > 0$). This is true for both altruist-cheater polymorphism (the area of such polymorphism is visible in yellow for a level of investment $I = 0$ on the X-axis or the Y-axis) or altruist-altruist polymorphism (yellow area everywhere else, i.e., where both levels of investment are greater than 0). The persistence of polymorphism between two altruist genotypes (or between a cheater and an altruist one) will ultimately depend on what levels of investment are biologically possible (i.e., what values could be shown by a mutant strategy): if the ESS (red star in Fig. S4) is not attainable and the two nearest feasible phenotypes lie on either side of it—and are sufficiently close to the ESS that each falls within the other's partial-invasion region—then a stable polymorphism emerges between a stingy and an overly generous altruist.

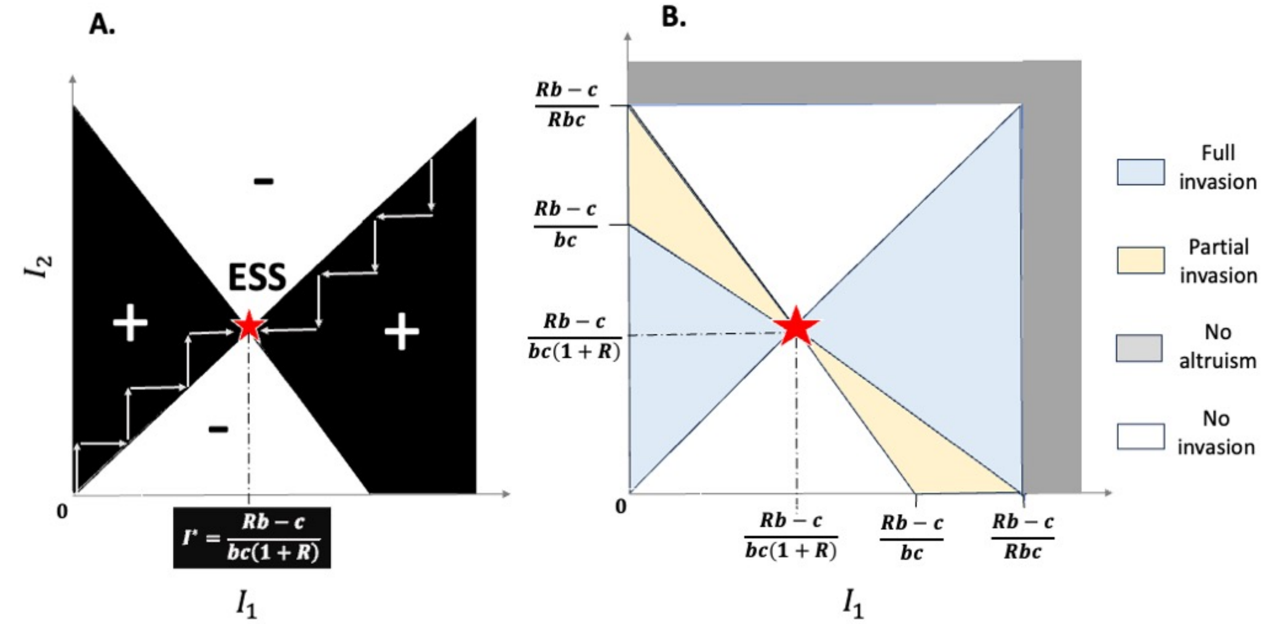


**Figure S4. Illustration of the Adaptive Dynamics analysis of the level of investment**. **A**) The pairwise invasibility plot shows the invasion fitness of a mutant allele that invests at a level $I_2$ of its resources into public goods, in a population where the resident allele invests $I_1$, where $I_1$ and $I_2$ can both be 0 (and hence could represent cheaters/non-altruists). When the invasion fitness of a mutant is positive (shown as a black area marked with a '+'), mutants invade and replace the resident strategy, whereas the resident strategy remains in place if the mutant invasion fitness is negative (shown as a white area marked with a '–'). The singular strategy, $I^* = (Rb - c)/((1 + R)bc)$ is an ESS (red star) because it cannot be invaded by mutants investing a lower or higher level of resources into public goods (which is demonstrated by the presence of '– ' areas above and below it on the figure). Mutants investing at the ESS level always invade other strategies (shown by the presence of the black '+' area on the left and the right sides of the ESS, all along the parameter space), making that strategy convergent and theoretically reachable because there always exists a large mutational range whose invasive fitness is positive. **B**) We delineate two different cases when rare mutants invade: in the blue shaded area, full invasion occurs (mutants cannot be invaded by the previous resident so that they go to fixation), while in the yellow shaded area, the previous resident can invade a population where the mutant becomes the new resident, which would therefore lead to a polymorphic equilibrium. At the level of the ESS (red star), full invasion by a mutant always occurs regardless of the investment level of the resident strategy ($I_1$). For a resident investing at the level of a cheater ($I_1 = 0$), mutants investing between $(Rb - c)/(bc)$ and $(Rb - c)/(Rbc)$ invade partially (i.e., they establish a polymorphism), because cheater mutants ($I_2 = 0$) can also invade partially resident strategies in between these two boundaries. The (yellow shaded) partial invasion area decreases as the resident strategy gets closer to the ESS ($I_1 \to (Rb - c)/[(1 + R)bc]$.

To determine whether or not a mutant closer to the ESS, denoted '$A_3$', can invade in these conditions, we need to evaluate whether it has a higher invasive fitness in the polymorphic environment set by the coexistence between '$A_1$' and '$A_2$' (where '$A_2$' has now a non-vanishing frequency $p_2 = 1 - p_1$ in the population corresponding to the evolutionary equilibrium). The respective fitness of '$A_1$' and '$A_2$' in a dimorphic environment $p_1 + p_2 \to 1$, subscripted $|d$, comprised of both of them is given by:

$$\begin{cases} \overline{W}_{A_1|d}(I_1; I_2) = (1 - cI_1)(1 + b(R + (1-R)p_1)I_1 + b(1-R)(1-p_1)I_2) \\ \overline{W}_{A_2|d}(I_2; I_1) = (1 - cI_2)(1 + b(R + (1-R)p_2)I_2 + b(1-R)(1-p_2)I_1) \end{cases} \quad \text{(S17)}$$

Considering that they have frequencies $p_1$ and $p_2$ gives the following fitness difference (similar to the equation on the evolutionary change, except for the genetic variance term, in the main text that leads to eqn. 12):

$$\Delta\overline{W}_{2-1} = (I_1 - I_2)\big(Rb - c - bcI_1(R + (1-R)p_1) - bcI_2(R + (1-R)p_2)\big) \quad \text{(S18)}$$

The equilibrium is reached when this difference (proportional to the selection coefficient) cancels out, which can be shown to coincide with the following condition:

$$p_1^* = \frac{Rb - c - bcRI_1 - bcI_2}{bc(I_1 - I_2)(1 - R)} \quad \text{(S19)}$$

This can also be written in the same form as equation (12). The fitness of each of these resident strategies at equilibrium is thus given by $\overline{W}^* = \overline{W}^*_{A_1|d} = \overline{W}^*_{A_2|d} = (1 - cI_1)(1 - cI_2)Rb/c$, while the fitness of a new mutant, denoted $A_3$, equals (after a few straightforward steps of calculus):

$$\overline{W}^*_{A_3|d} = (1 - cI_3)(1 + cI_3 - c(I_1 + I_2))Rb/c \quad \text{(S20)}$$

The conditions under which this third allele invades correspond to $\overline{W}^*_{A_3|d} > \overline{W}^*$. It is possible to write the selection difference of the third allele as:

$$\overline{W}^*_{A_3|d} - \overline{W}^* = -c^2(I_3 - I_1)(I_3 - I_2) \quad \text{(S21)}$$

This leads to the condition $(I_3 - I_1)(I_3 - I_2) < 0$ for $A_3$ to invade, implying that any mutant allele closer to the ESS has a higher fitness than the two coexisting resident alleles (because the factors require opposite signs for that condition to be verified). Furthermore, only the resident whose investment is on the opposite side than the mutant with regards to the ESS (for example, higher than the ESS if the third/invading allele is below the ESS) can invade back (see Fig. S4B), which implies that the third allele either goes to fixation or establishes a polymorphism with this latter by driving out the resident who shares its overgenerosity or stinginess.

#### Local competition scenario

We have shown above that local competition can alter the conditions that favour the altruist or the non-altruist genotype, which ends up reducing the parameter space conducive to polymorphism. Here, we show that local competition has no influence on the ESS other than limiting kin selection benefits to the component of relatedness that is competition independent (which represents local relatedness, corresponding to variation among groups within local areas; (Queller, 1994)).

To demonstrate that local competition does not alter the ESS, we first show that the singular strategy under local competition is equal to the singular strategy within local areas. Starting back from equation (S12), a strategy is singular under local competition if $\frac{\partial \overline{W}_{A_2}}{\partial I_2}|_{I_2 = I_1 = I^*_{comp}} = 0$ with $I^*_{comp}$ the population-level singular strategy. Although we derive results for the investment trait, the same argument applies to any trait under local competition, provided that the ESS outperforms neighbouring strategies at all mutant frequencies, a property that we prove explicitly below.

Let $\bar{W}_{A_2|p_{2(L)}}(I_2; I_1)$ denote the absolute fitness of a rare mutant allele $A_2$ in a local area with frequency $p_{2(L)}$ of this allele. Although the mutant has vanishing population frequency, it may experience all local area compositions. Under local competition (i.e., after rescaling by the local mean fitness), its global mean fitness is: $\bar{W}_{A_2|p_1\to 1} = \sum_{p_{2(L)}=0}^{1}\left(\bar{W}_{A_2|p_{2(L)}}/\bar{W}_{L|p_{2(L)}}\right)\ p_{2(L)}/p_1\ \ f\left(p_{2(L)}\right)$, where $\bar{W}_{L|p_{2(L)}}$ is the local mean fitness. Differentiating with respect to the mutant trait yields:

$$\frac{\partial \bar{W}_{A_2|p_1\to 1}}{\partial I_2} = \frac{\partial\left(\sum_{p_{2(L)}=0}^{1}\left(\frac{\bar{W}_{A_2|p_{2(L)}}}{\bar{W}_{L|p_{2(L)}}}\right)p_{2(L)}\ f\left(p_{2(L)}\right)\right)}{\partial I_2}$$
$$= \sum_{p_{2(L)}=0}^{1} \frac{\frac{\partial \bar{W}_{A_2|p_{2(L)}}}{\partial I_2}\bar{W}_{L|p_{2(L)}} - \bar{W}_{A_2|p_{2(L)}}\frac{\partial \bar{W}_{L|p_{2(L)}}}{\partial I_2}}{\bar{W}^2_{L|p_{2(L)}}} p_{2(L)} f\left(p_{2(L)}\right) \quad \text{(S22)}$$

We can then write the derivative close to the singular strategy under local competition ($I^*_{comp} = I_2 = I_1$), by noticing that this implies $\bar{W}_{L|p_{2(L)}} = \bar{W}_{A_2|p_{2(L)}} = (1 - cI^*_{comp})(1 + bI^*_{comp})$:

$$\frac{\partial \bar{W}_{A_2|p_1\to 1}}{\partial I_2}|_{I_2=I_1=I^*_{comp}} = \lim_{p_1\to 1}\sum_{p_{2(L)}=0}^{1}\frac{\frac{\partial \bar{W}_{A_2|p_{2(L)}}}{\partial I_2} - \frac{\partial \bar{W}_{L|p_{2(L)}}}{\partial I_2}}{\bar{W}_{L|p_{2(L)}}} p_{2(L)} f\left(p_{2(L)}\right)$$
$$= \lim_{p_1\to 1}\sum_{p_{2(L)}=0}^{1}\frac{\partial(\bar{W}_{A_2|p_{2(L)}} - \bar{W}_{L|p_{2(L)}})/\partial I_2}{\bar{W}_{L|p_{2(L)}}} p_{2(L)} f\left(p_{2(L)}\right) \quad \text{(S23)}$$

Equation (S23) shows that the global selection gradient is a weighted sum of local selection gradients, where we use selection gradients according to their meaning in adaptive dynamics. Now, we can show that the sign of these local gradients is identical in all local areas. Within a local area, we can write $\partial(\bar{W}_{A_2|p_{2(L)}} - \bar{W}_{L|p_{2(L)}})/\partial I_2 = (1 - cI^*_{comp})(1 + bI^*_{comp})\left(1 - p_{2(L)}\right)(R_{GL}b - c - bcI^*_{comp}(1 + R_{GL}))$ after a few steps of calculus. All terms but $(R_{GL}b - c - bcI^*_{comp}(1 + R_{GL}))$ are positive, $\partial \bar{W}_{A_2|p_1\to 1}/\partial I_2$ cancels out in all local areas when $I^*_{comp} = I^*_l = (R_{GL}b - c)/(1 + R_{GL})$, with $I^*_{comp}$ being a singular strategy. Moreover, the sign of the local gradient is the same across local areas (the local gradient is positive when $I < I^*_{comp}$, and vice versa). By consequence, the global selection gradient can only vanish when all its components vanish, and hence, $I^*_{comp}$ is the single singular strategy.

Finally, we verify uninvadability. From equation (12) in the main text, the mutant-resident fitness difference near the local ESS can be written as $\bar{W}\Delta p_{2(L)} = \left[Rb - c - bc\left(\left(R + (1 - R)p_{2(L)}\right)\delta + (1 + R)I^*_{comp}\right)\right]$ for a small deviation $\delta = I_2 - I^*_{comp}$ (note that indices are different as $I_1 = I_2 + \delta$ in the main text). Plugging $I^*_{comp}$ in this expression, we can write $\bar{W}\Delta p_{2(L)} = -\delta(R + (1 - R)p_{2(L)})$, which is strictly negative and confirms that $I^*_{comp}$ is the ESS under local competition. Since $\frac{\partial \bar{W}_{A_2|p_1\to 1}}{\partial I_2}|_{I_2=I_1=I} > 0$, when $I < I^*_{comp}$, and negative otherwise, the singular strategy also invades neighbour residents, making $I^*_{comp}$ convergent stable, and more generally, a CSS under local competition.

#### Supporting Text S4: Evolutionary dynamics with local competition

To understand the impact of local competition, which imposes a constraint because all local areas have the same mean fitness, we can first reconsider the case in the absence of local competition. We could calculate the change in the frequency of the altruist allele within each local area by first calculating the local selection coefficient, $s_{A|p_{A(L)}}$, which represents the difference in the relative fitness of the two alleles within a local area that has a $p_{A(L)}$ frequency of altruists: $s_{A|p_{A(L)}} = \left(\overline{W}_{A|p_{A(L)}} - \overline{W}_{N|p_{A(L)}}\right)/\overline{W}_{L|p_{A(L)}}$, where $\overline{W}_{A|p_{A(L)}}$ and $\overline{W}_{N|p_{A(L)}}$ are the average fitnesses of the two alleles within that level and $\overline{W}_{L|p_{A(L)}}$ is the average fitness of that local area. The local change in the frequency of the altruist is then simply the local selection coefficient times the local allelic variance: $\Delta p_{A(L)} = s_{A|p_{A(L)}} p_{A(L)} p_{N(L)}$ where $p_{A(L)}$ and $p_{N(L)}$ are the local frequencies of the altruist and non-altruist alleles respectively. Weighting these local changes by the relative mean local fitness, $\overline{W}_{L|p_{A(L)}}/\overline{W}$, and summing over all local areas returns the global change (since the local mean fitness term in the local change cancels out with the weighting by relative local mean fitness):

$$\begin{aligned}\Delta p_{A(total)} &= \sum_{p_{A(L)}=0}^{1} \frac{\left(\overline{W}_{A|p_{A(L)}} - \overline{W}_{N|p_{A(L)}}\right)}{\overline{W}_{L|p_{A(L)}}} p_{A(L)} p_{N(L)} \frac{\overline{W}_{L|p_{A(L)}}}{\overline{W}} f\left(p_{A(L)}\right) \\ &= \frac{\overline{W}_{A|p_A} - \overline{W}_{N|p_A}}{\overline{W}} p_A p_N \\ &= s_A p_A p_N\end{aligned} \tag{S24}$$

When there is a competitive constraint that causes all local areas to have the same mean fitness, this simply alters the summation in equation (S23) such that local areas are all weighted equally (since they all have the same relative fitness) instead of being weighted by their local relative fitness:

$$\Delta p_{A(competition)} = \sum_{p_{A(L)}=0}^{1} \left(\frac{\overline{W}_{A|p_{A(L)}} - \overline{W}_{\mathrm{N}|p_{A(L)}}}{\overline{W}_{L|p_{A(L)}}}\right) p_{A(L)} p_{N(L)}\, f\left(p_{A(L)}\right) \tag{S25}$$

which recovers the intermediate form of equation (18).

Equation (S25) can be simplified by using the zeroth-order approximation for the average of a ratio, which is given by the ratio between the average of the numerator and that of the denominator:

$$\Delta p_{A(competition)} \approx \frac{\sum_{p_{A(L)}=0}^{1} \left(\overline{W}_{A|p_{A(L)}} - \overline{W}_{\mathrm{N}|p_{A(L)}}\right) p_{A(L)} p_{N(L)}\ f\left(p_{A(L)}\right)}{\sum_{p_{A(L)}=0}^{1} \overline{W}_{L|p_{A(L)}}\, f\left(p_{A(L)}\right)} \tag{S26}$$

Because the denominator is constrained to be greater than zero, the sign of the evolutionary change given by (S26) depends solely on the numerator. This implies that the evolutionary outcome given by (S26), in terms of whether the change in the frequency of the altruist allele is positive, negative, or zero, will be identical if one neglects the denominator to calculate $\Delta p_{A(competition)}$, which explains why making this assumption (as we do in the main document; see eqn. 18) gives a relatively accurate approximation for equilibrium. Moreover, we can rewrite the approximation in (S26) by considering that, under relatively weak selection, $\overline{W}_{L|p_{A(L)}} \approx 1$, which gives:

$$\Delta p_{A(competition)} \approx \sum_{p_{A(L)}=0}^{1} \left(\bar{W}_{A|p_{A(L)}} - \bar{W}_{\mathrm{N}|p_{A(L)}}\right) p_{A(L)} p_{N(L)}\ f\left(p_{A(L)}\right) \tag{S27}$$

Since mean fitness can be written as a function of investment $\bar{W}_{L|p_{A(L)}} = 1 + (b-c)I\,\hat{p}_A + o(I)$ for both the trade-off and additive scenarios, neglecting the denominator works better when $I \ll 1$ and is hence akin to the kin theory version of weak selection (Wild & Traulsen, 2007).

For the rate of evolutionary change, approximating the average of a ratio by the ratio of the average introduces a systematic bias that tends to increase the result, given that $\sum_{p_{A(L)}=0}^{1}\left(\bar{W}_{L|p_{A(L)}}\right) p_{A(L)} p_{N(L)}\, f\left(p_{A(L)}\right) > 1$ for a wide range of the parameter space (wherever cooperation enhances group fitness; for example, when $b > c$ in the additive case). For these reasons, the approximate evolutionary change usually also represents an upper bound of the exact quantity.

We next derive the exact solution for equation (19) under the additive scenario, which provides a guide as to where approximations might be inaccurate. After that, we provide further explanations for the derivation of the approximations for the trade-off case (eqns. 20-22) and contrast the results of the analytical approximation with numerical solutions.

#### Additive costs and benefits

In order to solve equation (18) for the additive case, we first need to calculate the difference in local fitness between the altruist and the non-altruist alleles:

$$\frac{\bar{W}_{A|p_{A(L)}} - \bar{W}_{N|p_{A(L)}}}{\bar{W}_{L|p_{A(L)}}} = \frac{(R_{GL}b - c)I}{1 + (b-c)Ip_{A(L)}} \tag{S28}$$

We can then substitute this expression into equation (18), which gives:

$$\Delta p_{A(competition)} = \sum_{p_{A(L)}=0}^{1} \frac{(R_{GL}b - c)I}{1 + (b-c)Ip_{A(L)}} p_{A(L)} p_{N(L)} f\left(p_{A(L)}\right) \tag{S29}$$

We can remove the term in the numerator that corresponds to the fitness differential in Hamilton's rule from the summation since it does not depend on allele frequencies:

$$\Delta p_{A(competition)} = (R_{GL}b - c)I \sum_{p_{A(L)}=0}^{1} \frac{p_{A(L)} p_{N(L)}}{1 + (b-c)Ip_{A(L)}} f\left(p_{A(L)}\right) \tag{S30}$$

If we assume that $f\left(p_{A(L)}\right)$ coincides here with the density function of a Beta distribution (so we are neglecting the finite group size effect, which is akin to assuming that local areas population sizes are large), the full expression can be written as :

$$\Delta p_{A(competition)} = \frac{(R_{GL}b - c)I}{B(\theta_L p_A, \theta_L p_N)} \int_{(p_{A(L)}=0}^{1} \frac{(p_{A(L)})^{\theta_L p_A}\, p_{N(L)}{}^{\theta_L p_N}}{\left(1 + (b-c)Ip_{A(L)}\right)} dp_{A(L)} \tag{S31}$$

where $\theta_L = \frac{1}{R_{LT}} - 1$. The solution to the integral can be expressed as a hypergeometric series using the Euler integral formula:

$$\Delta p_{A(competition)} = (R_{GL}b - c)I(1 - R_{LT})p_A p_N \ {}_2F_1[j,k;l;z] \tag{S32}$$

where ${}_2F_1[j,k;l;z]$ represents the ordinary hypergeometric function, and whose parameters are given in our case by $j = 1$, $k = 1 - \left(1 - \frac{1}{R_{LT}}\right) p_A$, $l = 1 + 1/R_{LT}$ and $z = -(b-c)I$ . Considering small investment values $I \ll 1$, we can expand equation (S32) using the power series definition of the hypergeometric function:

$$\Delta p_{A(competition)} = (R_{GL}b - cI)(1 - R_{LT})p_A p_N \times (1 + H(1;R_{LT})[(b-c)I] + o[(b-c)I]) \tag{S33}$$

where $H(1;R_{LT}) = -(R_{LT} + (1 - R_{LT})p_A)/(R_{LT} + 1)$. Using this equation, we can recover equation (19) as a zeroth-order approximation and we can notice that this approximation should perform well when $(b-c)I$ is low, and even more so when either or both $R_{LT}$ and $p_A$ are relatively low. More generally, the full numerical series, whose higher order terms are neglected in equation (S33), is convergent provided $(b-c)I < 1$ , which corresponds to the weak selection approximation when $I$ is close to zero. Under the weak selection scenario, the trade-off case (which we explain below) is similar to the additive scenario as the opportunity cost scales with $I^2$ and is no longer relevant: hence, the additive scenario explains why we do not observe polymorphism in the trade-off case for low investment values (see Fig. S5). Note finally that if we use the method described in the main document, meaning we use equation (S27) and thus neglect the denominator in equation (S30), we directly recover the zeroth-order approximation as:

$$\Delta p_{A(competition)} \approx (R_{GL}b - c)I \sum_{p_{A(L)}=0}^{1} p_{A(L)} p_{N(L)} f\left(p_{A(L)}\right) \tag{S34}$$

where $\sum_{p_{A(L)}=0}^{1} p_{A(L)} p_{N(L)} f\left(p_{A(L)}\right)$ is the genetic variance among local areas.

#### Resource trade-off model

Under the trade-off scenario, the opportunity cost introduces a quadratic effect of the allelic frequency on local mean fitness, and hence, in the denominator of equation (S25):

$$\frac{\overline{W}_{A|p_{A(L)}} - \overline{W}_{N|p_{A(L)}}}{\overline{W}_{L|p_{A(L)}}} = \frac{(R_{GL}b - c)I - bcI^2(R_{GL} + (1 - R_{GL})p_{A(L)})}{1 + (b-c)Ip_{A(L)} - bcI^2 p_{A(L)}(R_{GL} + (1 - R_{GL})p_{A(L)})} \tag{S35}$$

Because of the quadratic effect in the denominator, the exact solution to the total evolutionary change becomes more convoluted. It is nonetheless possible to find the simple zeroth-order approximation that focuses on the integration of the numerator:

$$\begin{aligned} \Delta p_{A(competition)} &= \sum_{p_{A(L)}=0}^{1} \frac{(R_{GL}b - c)I - bcI^2(R_{GL} + (1 - R_{GL})p_{A(L)})}{1 + (b-c)Ip_{A(L)} - bcI^2 p_{A(L)}(R_{GL} + (1 - R_{GL})p_{A(L)})} p_{A(L)} p_{N(L)} \, f\left(p_{A(L)}\right) \\ &\approx \sum_{p_{A(L)}=0}^{1} (R_{GL}b - c)I - bcI^2(R_{GL} + (1 - R_{GL})p_{A(L)}) \, p_{A(L)} p_{N(L)} \, f\left(p_{A(L)}\right) \end{aligned} \tag{S36}$$

The result of the part involving Hamilton's fitness differential is the same as in the additive case, so that this expression directly yields equation (20).

In order to understand the effect of the different levels of relatedness, and the benefit-to-cost ratio on the evolutionary outcome, we can first plot the case where the investment level is very low, which comes back to the traditional additive outcome where altruism either gets fixed (if Hamilton's rule is satisfied within local areas) or does not invade at all (otherwise), which is shown in Figure S5. Note that all figures generated in the supplements are based on a script that is available on GitHub (https://github.com/FloTuzoLab/Evolution-public-goods-altruism).

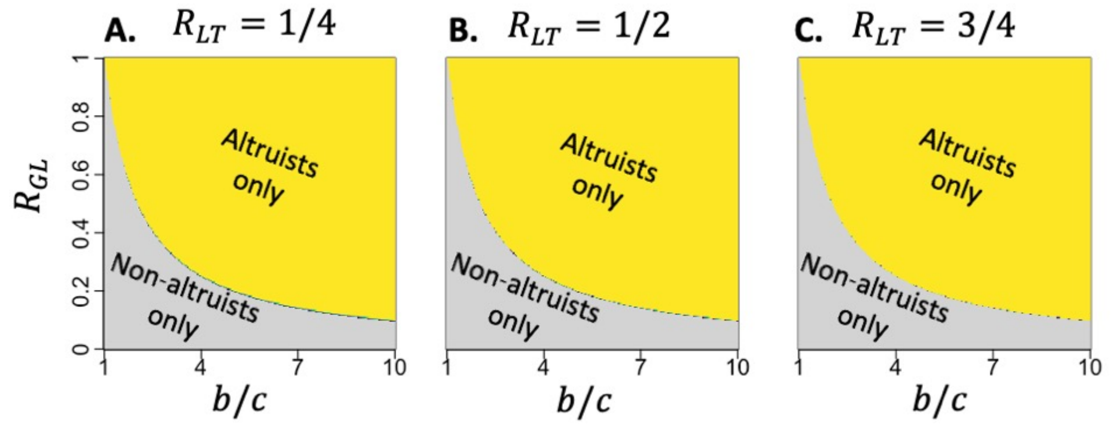


**Figure S5. Evolutionary outcome for the trade-off scenario when investment level is very low ($I \approx 0$).** Areas in each plot are shaded according to which allele is favoured as a function of $b/c$ and $R_{GL}$. Regions where altruists are favoured appear in yellow, while areas where non-altruists are favoured appear in grey. When investment is infinitesimal, there is effectively no region that supports polymorphism, so the boundary between the two outcomes is given by Hamilton's rule, $R_{GL}b - c > 0$. Note that relatedness at the level of local areas, $R_{LT}$ (increasing from **A** to **C**) has no influence on the outcome because any of the kin-selected benefit it provides is cancelled out by extra local competition.

To visualise the influence of local competition, it is useful to first visualise the outcomes under the simplest case of global competition (that coincides with $R_{LT} \to 0$), for which the analytical results are given by equations (8-9) (see Fig. S6).

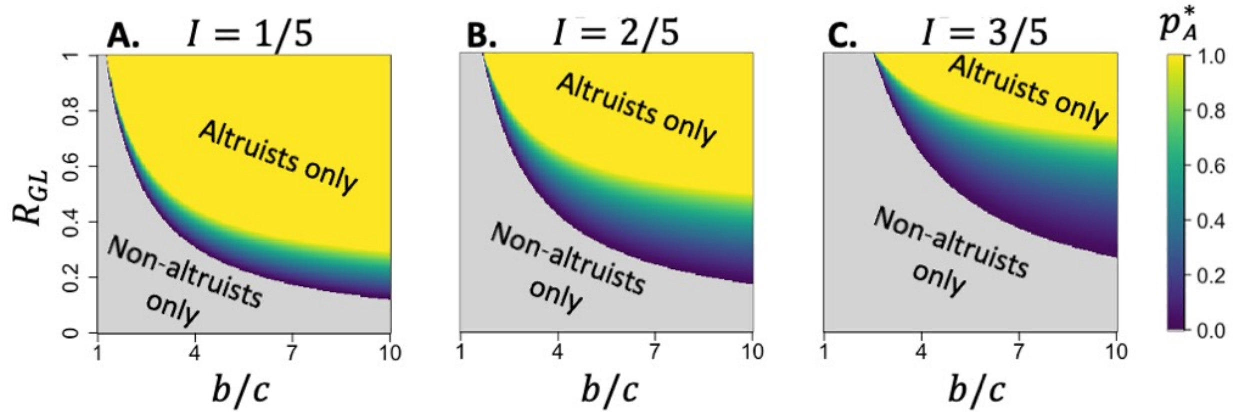


**Figure S6. Evolutionary outcome for the trade-off scenario under global competition ($R_{LT} = 0$).** Areas in each plot are shaded according to which allele is favoured as a function of $b/c$ and $R_{GL}$. Regions where altruists are favoured appear in yellow, areas where non-altruists are favoured appear in grey, and regions where there is polymorphism are shaded along a blue-green-yellow continuum that corresponds to the equilibrium frequency of altruists (see the legend to the RHS). The parameter space supporting polymorphism increases as the investment level, $I$, gets larger (from **A** to **C**)

To show how local competition impacts the evolutionary outcome between altruists and non-altruists (i.e., whether either is universally favoured, or the conditions allow for polymorphism), we can visualise how the outcome is jointly influenced by the component of relatedness that corresponds to variation in allele frequencies across local areas ($R_{LT}$) and the level of investment (see Fig. S7). As described in the manuscript, local competition decreases the parameter space where altruists are favoured and increases the parameter space where non-altruists (cheaters) are favoured. Together, this ends up reducing the parameter space that allows for polymorphism (see Fig. 2). Overall, we see that the parameter space supporting polymorphism is highest when $R_{GL}$ is intermediate, benefits are high relative to costs, local competition is low (low $R_{LT}$), and investment levels $I$ are relatively high (see Figs. S6 and S7).

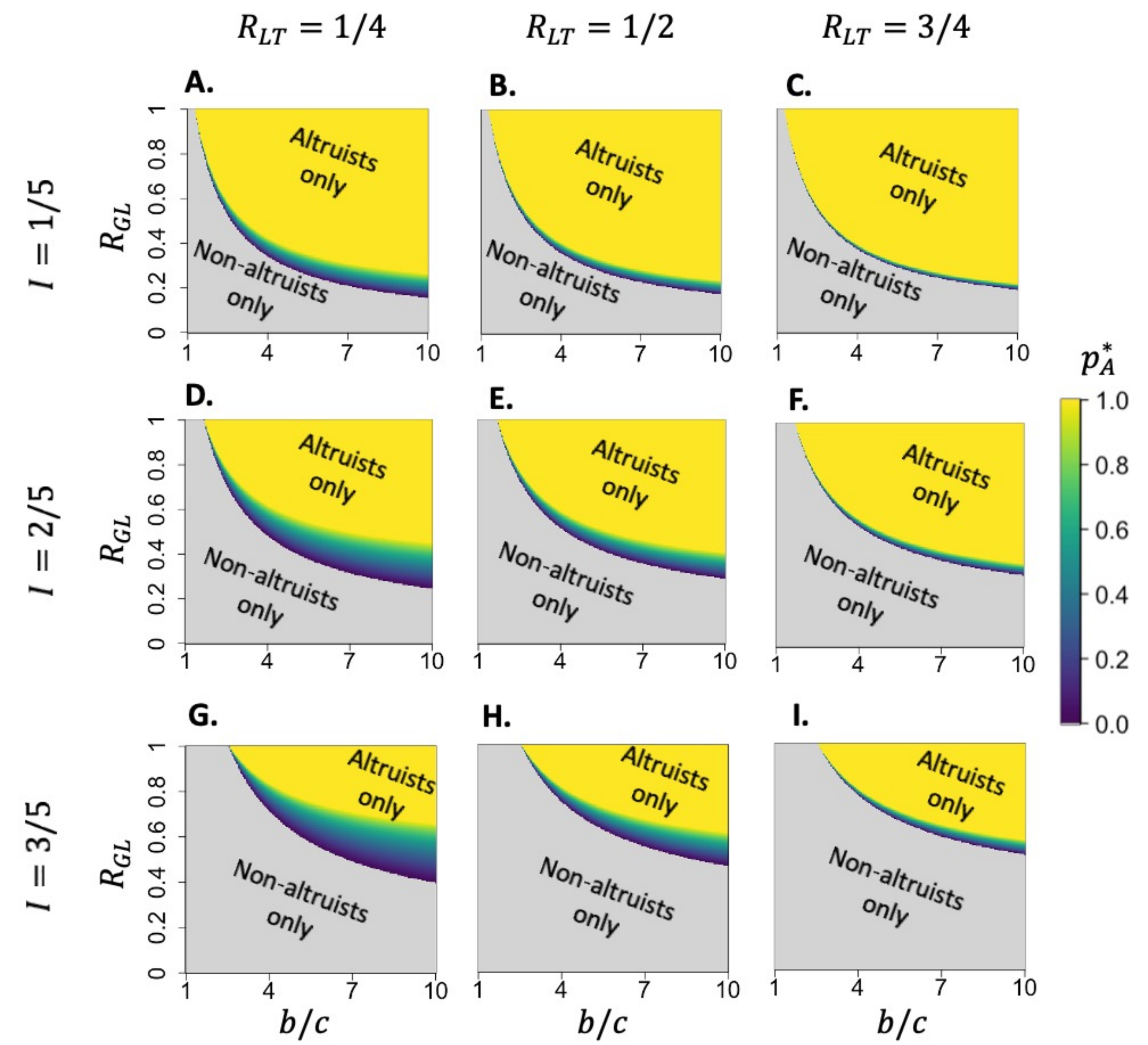


**Figure S7. Evolutionary outcome for the trade-off scenario under local competition ($R_{LT} \neq 0$).** Areas in each plot are shaded according to which allele is favoured as a function of $b/c$ and $R_{GL}$. Regions where altruists are favoured appear in yellow, areas where non-altruists are favoured appear in grey, and regions where there is polymorphism are shaded along a blue-green-yellow continuum that corresponds to the equilibrium frequency of altruists (see the legend to the RHS). The outcomes are shown in terms of the level of local competition (increasing cross columns from left to right) and the level of investment (increasing in rows from top to bottom).

#### Accuracy of approximations for the resource trade-off model

To evaluate the performance of the zeroth-order approximations we can compare them against the numerical solutions. In general, the approximation is slightly more stringent (in favour of non-altruists), especially with respect to the boundary for the emergence of polymorphism, with the thresholds being underestimated by a few to ten percent across a wide range of parameters (see Fig. S8). As expected (from the comparison between the exact and approximate solutions in the additive case), the approximation loses accuracy as $I$ increases (so that the approximations are slightly closer to the exact values for fixation than for polymorphism), when local competition $R_{LT}$ and $b/c$ gets larger, and when $R_{GL}$ gets lower. This implies that altruists might still be favoured even when investment levels are slightly higher than the approximation for the value below which altruists would be universally favoured, $I_{altruist} = \frac{(R_{GL}b-c)(1+R_{LT})}{bc\,(1+R_{LT}R_{GL})}$ (see Fig. S8A to C). Likewise, polymorphism may be possible for investment levels higher than the approximation for the value above which cheaters would be universally favoured, $I_{cheater} = \frac{(R_{GL}b-c)(1+R_{LT})}{bc\,(R_{LT}+R_{GL})}$ (see Fig. S8D to F). More generally, the approximation slightly underestimates the proportion of altruists at the evolutionary equilibrium.

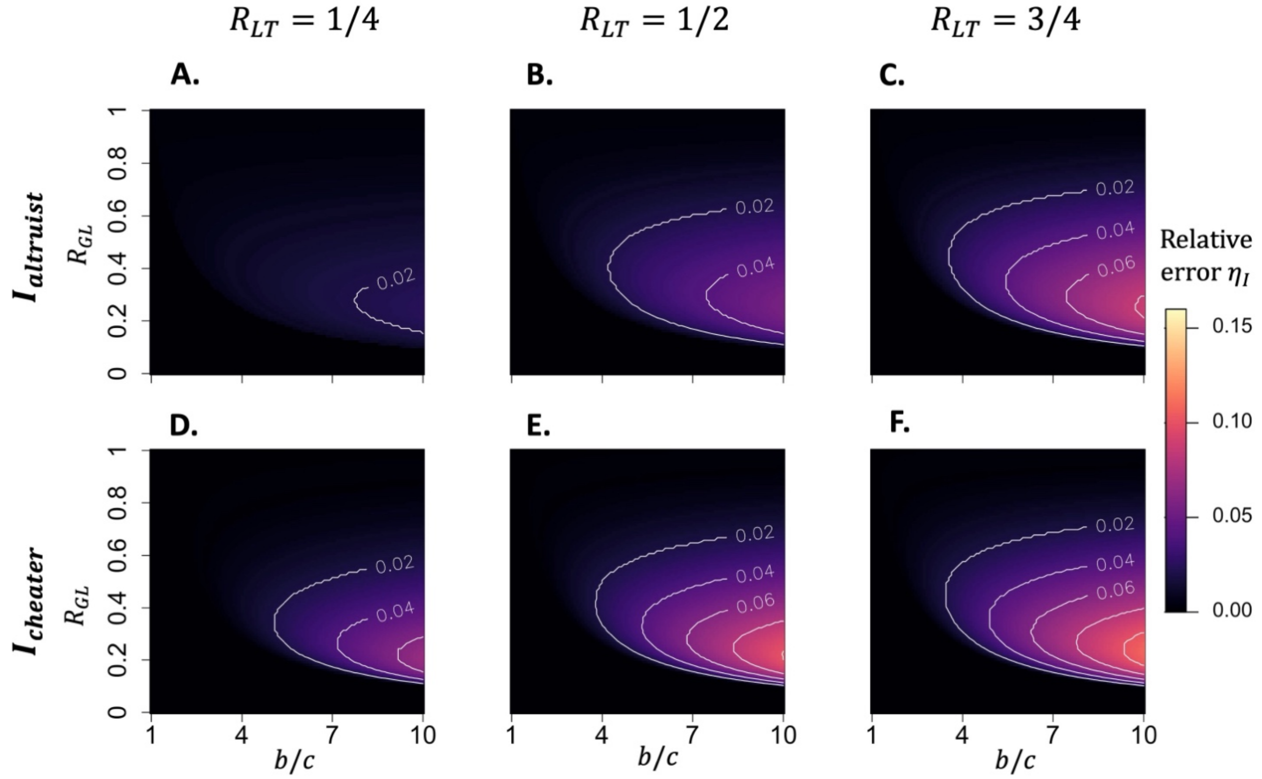


**Figure S8. Comparison of the evolutionary outcomes for the zeroth-order approximation (ratio of averages) and the exact numerical solution.** Each panel shows the relative error of the approximate solution compared to the exact numerical solution, calculated as $\eta_I = (I_{numerical} - I_{approximation})/I_{numerical}$, represented by the 'elevation' of the contours (with lighter colours denoting lower accuracy; see the legend on the RHS) as a function of the benefit/cost ratio ($b/c$) and level of local relatedness (which represents the level of variation among groups within local areas), $R_{GL}$. Note that all error values are positive, which indicates that the approximation always underestimates the thresholds. The relative error ($\eta_I$) in the approximation for the level of investment ($I_{altruist}$) below which the altruist allele is favoured (first row, **A** to **C**) and the level of investment ($I_{cheater}$) above which cheaters are universally favoured (second row, **D** to **F**) are shown for different levels of variation among local areas, $R_{LT}$ (columns).

#### Supporting Text S5: Game-theoretical perspective on the evolution of publics good altruism

In evolutionary game theory, different games (or rules for the game) can be classified according to their expected evolutionary outcomes (Doebeli & Hauert, 2005; J. Maynard Smith & Price, 1973). Here, we exploit this logic to relate our framework to well-known games and to show how our model can be interpreted as a generalisation of these games, where the players in the game are the alternative alleles. In typical evolutionary games, the payoff matrix describes the outcome of an encounter between a pair of individuals who each adopt a particular strategy (Maynard Smith, 1982). However, for public goods games, interactions are not pairwise, and payoffs depend on the total level of public goods investment within groups of given sizes. Therefore, we adopt an approach where payoffs are defined for a rare focal allele in a population that is effectively composed entirely of a resident allele (as done in adaptive dynamics). We show a generic pay-off matrix for altruist-cheater (i.e., non-altruist) games in Table S1, which describes the gain (in terms of fitness) that the focal allele (in the row) receives when it is in a population comprised of alleles playing the resident strategy (in columns). The payoffs are denoted $P_{XY}$, with the first subscript (with $A$ = 'altruist' and $C$ = 'cheater'/'non-altruist') giving the focal allele and the second the resident. When an allele plays against itself as the resident allele, both players have the same fitness.

**Table S1**. **A generic payoff matrix for altruist-cheater games.** The rows correspond to the focal strategy (altruist or cheater) that is in a population composed entirely of the resident strategy in the columns. The payoffs are denoted $P_{XY}$, with the first subscript being the focal strategy and the second the resident.

| **Resident→ Focal↓** | **Altruist** | **Cheater** |
|---|---|---|
| **Altruist** | $P_{AA}$ | $P_{AC}$ |
| **Cheater** | $P_{CA}$ | $P_{CC}$ |

By considering the fitness of a rare focal allele, the payoffs can be compared to evaluate whether the focal allele would invade a population composed of the resident allele. For example, if $P_{CA} > P_{AA}$ then the fitness of cheaters is higher than that of altruists when the population is almost entirely comprised of altruists. Consequently, cheaters would invade. The qualitative outcome (whether one allele would go to fixation) would then simply depend on the ranking of payoffs.

Social dilemmas arise when a population of altruists does better than a population of cheaters, i.e., $P_{AA} > P_{CC}$, but the free-rider benefit that cheaters get in a population of altruists prevents altruism fixation. Two such dilemmas have been proposed depending on their outcome. Firstly, in the prisoner's dilemma (PD), cheaters go to fixation because the ordering of the payoffs is $P_{CA} > P_{AA} > P_{CC} > P_{AC}$. Moreover, because $P_{CC} > P_{AC}$, altruists cannot even invade (when being rare). Secondly, in the snowdrift game (SD), polymorphism can be maintained because both genotypes can invade in a resident population of the other one, which arises because the ordering of the payoffs is $P_{CA} > P_{AA} > P_{AC} > P_{CC}$. This phenomenon explains why the emergence of cheater-altruist polymorphism has been linked to SD scenarios.

#### Public goods games with additive costs and benefits and the role of kin selection

To consider the influence of kin selection, payoffs can be calculated based on inclusive fitness. For this we can first consider a very simple game that follows the logic of the scenario with additive costs and benefits that we present in the main text (where altruism provides a benefit, $b$, and altruists pay a cost, $c$). The payoffs for this game are shown in Table S2.

**Table S2. The payoff matrix for a public goods game with additive costs and benefits.** The resulting payoff structure follows that of the prisoner's dilemma. The structure of the payoff matrix is described in the legend for Table S1.

| **Resident→ Focal↓** | **Altruist** | **Cheater** |
|---|---|---|
| **Altruist** | $b - c$ | $-c$ |
| **Cheater** | $b$ | $0$ |

Assuming that the benefits of altruism are larger than the costs (which is obviously a necessary condition for altruism to be favoured under any conditions), the ranking of the payoffs in this game match the PD, where $b > b - c > 0 > -c$. This game can be converted into the SD by assuming that altruists receive a proportion of the benefit even in a population of cheaters (as seen in some public goods games). In this case, the payoff to an altruist in a population of cheaters could be $P_{AC} = \gamma b - c$, which can be greater than $P_{CC}$ provided the value of $\gamma$ and the ratio $b/c$ are large enough. A simple way for this scenario to be true for the case of public goods is for group sizes to be small, such that an individual altruist necessarily receives a share of the benefits. This effect is captured by the $1/G$ component of whole-group relatedness (eqn. 4). To consider kin selection more broadly (i.e., beyond the simple benefits to self), we can modify the payoffs in Table S2 to account for relatedness. For this, we first calculate the average fitness of each allele across a population of groups, which matches the solutions for the values of $\overline{W}_A$ and $\overline{W}_N$ for the scenario of additive costs and benefits. We then derive the solution to these expectations for the condition where the frequency of the focal allele approaches zero. The payoffs under this scenario are shown in Table S3. Relatedness alters the payoffs in the cheater-altruist combinations ($P_{AC}$ and $P_{CA}$) from those in Table S2 because relatedness biases interactions towards self.

**Table S3. The payoff matrix for a public goods game with additive costs and benefits and relatedness.** The structure of the payoff matrix is described in the legend for Table S1.

| **Resident→ Focal↓** | **Altruist** | **Cheater** |
|---|---|---|
| **Altruist** | $b - c$ | $Rb - c$ |
| **Cheater** | $(1 - R)b$ | $0$ |

The payoffs return Hamilton's rule for the invasion and fixation of the altruist allele. When relatedness is low the game approaches the PD conditions for the game in Table S2, but once Hamilton's rule is met, the ESS strategy is altruism because these conditions mean that $P_{AA} > P_{CA}$ and $P_{AC} > P_{CC}$.

#### Public goods games with resource allocation trade-offs

Following our approach in the main text, we can modify the additive public goods game above by adding the cost of resource allocation trade-offs. This produces a payoff matrix similar to that in Table S3, but with the addition of the opportunity cost term (see Table S4).

**Table S4. The payoff matrix for a public goods game with resource allocation trade-offs.** The structure of the payoff matrix is described in the legend for Table S1.

| Resident→ Focal↓ | Altruist | Cheater |
|---|---|---|
| **Altruist** | $b - c - bcI$ | $Rb - c - RbcI$ |
| **Cheater** | $(1 - R)b$ | $0$ |

Using this payoff matrix, we can identify the conditions where the pattern of payoffs (in terms of their rank order) match each of the games described above. We can identify the conditions that universally favour altruism by evaluating the conditions where $P_{AA} > P_{CA}$ and $P_{AC} > P_{CC}$, which returns $I < (Rb - c)/bc$. The evolutionary outcome matches that for the case of additive costs and benefits with relatedness shown in Table S3. This result emphasises the fact that the conditions for Hamilton's rule can be considered as a weak selection outcome of the case with trade-offs. The conditions corresponding to the SD game can be derived by identifying the parameter space (in terms of the level of investment) where the rank ordering of the payoffs is $P_{CA} > P_{AA} > P_{AC} > P_{CC}$. This produces the conditions $(Rb - c)/bc < I < (Rb - c)/Rbc$, which recovers our finding for the conditions required for there to be polymorphism. Finally, we can follow this same approach to identify the conditions where the ordering of the payoffs matches the PD: $P_{CA} > P_{AA} > P_{CC} > P_{AC}$. This produces the condition $I > (Rb - c)/Rbc$, which matches the conditions we identified for where altruism cannot invade.

To understand the relationship between the public goods game with resource trade-offs and the classic evolutionary games (SD and PD), we can characterise how the evolutionary outcome of the game, in terms of $p_A^*$, changes with the level of investment $I$. In the main text (eqn. 9) we show that $p_A^* = (Rb - c - bcIR)/(bcI(1 - R))$. Using this equation, it can be shown that:

$$\frac{\partial p_A^*}{\partial I} = -K\,(Rb - c) \qquad \text{(S37)}$$

where $K > 0$. This relationship implies that the frequency of altruists at the evolutionary equilibrium decreases monotonically with the level of investment, meaning that increasing investment makes a system less and less favourable to altruism. This result also implies that the public goods game with tradeoffs shows phase transitions occurring when $p_A^* = 1$ (into SD) and when $p_A^* = 0$ (into PD). The sequence of equilibrium could be seen as 'monotonous' since the state variable $p_A^*$ always changes in the same direction when investment increases.
